# Two sites in the C-terminal β-chain tail mediate interactions of the chaperone clusterin with amyloid beta and other misfolded client proteins

**DOI:** 10.1101/2025.03.16.643389

**Authors:** Sandeep Satapathy, Ye Shen, Emma Jayne Proctor, Pietro Sormanni, Michele Vendruscolo, Mark R Wilson

**Author notes:** Corresponding author. Postal address: Rm 313, Building 42 (Molecular Horizons), University of Wollongong. Northfields Avenue, Wollongong. NSW. 2522. Australia;.

## Abstract

Clusterin (CLU) is a constitutively secreted mammalian chaperone that binds in extracellular body fluids to misfolded client proteins to neutralise their toxicity and mediate their safe disposal by cell uptake and intracellular degradation. However, the regions of CLU critical for its interactions with misfolded proteins remain still largely unknown. To identify binding sites, we expressed a panel of CLU deletion and alanine-stretch mutants in a recently developed mammalian expression system. Mutant CLU molecules lacking detectable structural aberrations were subjected to functional analyses to compare their abilities with that of wild type CLU to bind to misfolded proteins and to inhibit protein aggregation. These analyses implicated two regions in the flexible β-chain C-terminal tail of CLU as being important in the interactions of the chaperone with misfolded proteins, including aggregating the Alzheimer’s amyloid β-peptide (Aβ). We then designed *in silico* sequence-specific single-domain camelid nanobodies to confirm the function of the two putative client protein binding sites. Based on our experimental results and *in silico* binding site predictions, we suggest that the potent ability of CLU to promiscuously interact with many different misfolded proteins, regardless of their size or structure, arises from the location of multiple client protein binding sites in its flexible tail region.

## Introduction

Clusterin (CLU) is an extensively studied constitutively secreted mammalian chaperone known for its roles in both intra- and extra-cellular proteostasis [1]. In extracellular body fluids, CLU performs a critical function by binding to misfolded (client) proteins to neutralise their toxicity and by binding to cell surface receptors to mediate the uptake and intracellular degradation of CLU-client protein complexes within lysosomes [2]. This chaperone function protects the body from many different serious human diseases which are associated with the excessive accumulation of toxic misfolded proteins (e.g. Alzheimer’s and prion diseases, Type II diabetes) [1]. CLU has also been implicated in a variety of other important processes, including complement activation [3], cell proliferation [4], and cell-cell communication [5].

The structure of CLU is complex – the mature secreted molecule is an extensively glycosylated heterodimer, comprised of two chains (α- and β-) linked by 5 disulfide bonds. The conjugated sugars can comprise up to 30% of the total mass of CLU [6]. In physiological buffers, the CLU heterodimer associates to form a heterogenous mixture of differently sized oligomers [7]. Owing in large part to its complexity, there are currently no experimentally-determined structures for CLU. However, AlphaFold [8] has recently enabled confident predictions of at least large parts of CLU, which incorporates a rigid disulfide-bonded head region and four flexible α-helix-rich tails.

Despite its prominent roles in protein homeostasis processes critical to the support of life that occur both inside and outside cells [1], none of the regions of CLU important in binding to misfolded protein clients, cell receptors or its other important ligands have not yet been identified. CLU has limited sequence homology with any other known molecular chaperone, making it difficult to identify any misfolded protein binding sites based on sequence comparisons. To address this problem, we used mutagenesis as a primary tool to begin searching for CLU client binding sites. Initially, we constructed a series of CLU deletion mutants in which stretches of 30-37 residues were sequentially deleted from the full-length 449-residue molecule. These were expressed using a high-yield mammalian cell expression system for CLU we recently developed [9].

Those deletion mutants not suffering major structural aberrations were then compared with wild type CLU (^wt^CLU) for their ability to interact with Aβ and misfolded proteins. This identified the C-terminal region of the β-chain as likely to contain binding sites of interest. This region was subsequently more closely interrogated using scanning alanine-stretch mutagenesis, substituting short 8/9-mer stretches of alanine residues along parts of the β-chain and assessing the effects of these mutations on the interaction of CLU with Aβ and misfolded proteins. Through the combination of these approaches, we identified two regions in the CLU β-chain that putatively interact with both Aβ and misfolded proteins. We next used single-domain camelid nanobodies (dNbs) designed *in silico* to target specific CLU epitopes to confirm the function of the two client protein binding sites. Finally, incorporating AlphaFold predictions of the CLU structure, knowledge of the newly identified client binding sites, and other *in silico* predicted ligand binding sites, we propose a hypothetical molecular-level structure-function model for how CLU performs its life-supporting chaperone roles.

## Results

### Structure-function screening of CLU mutants

Before testing any CLU mutants in functional assays to identify specific regions of interest implicated in binding to client proteins, we first excluded any mutant associated with significant structural aberrations (Figure 1A). Owing to its extensive glycosylation, many regions of flexibility, and polydispersity in solution, CLU is challenging for traditional structural assays such as mass spectrometry and NMR spectroscopy. We therefore compared purified CLU mutants with ^wt^CLU in tractable structural assays, including SDS PAGE, immunoblotting, far-UV circular dichroism (CD) spectroscopy and far-UV melting curve analyses. Subsequent functional analyses allowed identification of CLU mutants that were structurally similar to ^wt^CLU but which had an impaired ability to interact with misfolded proteins.

**Figure 1.**
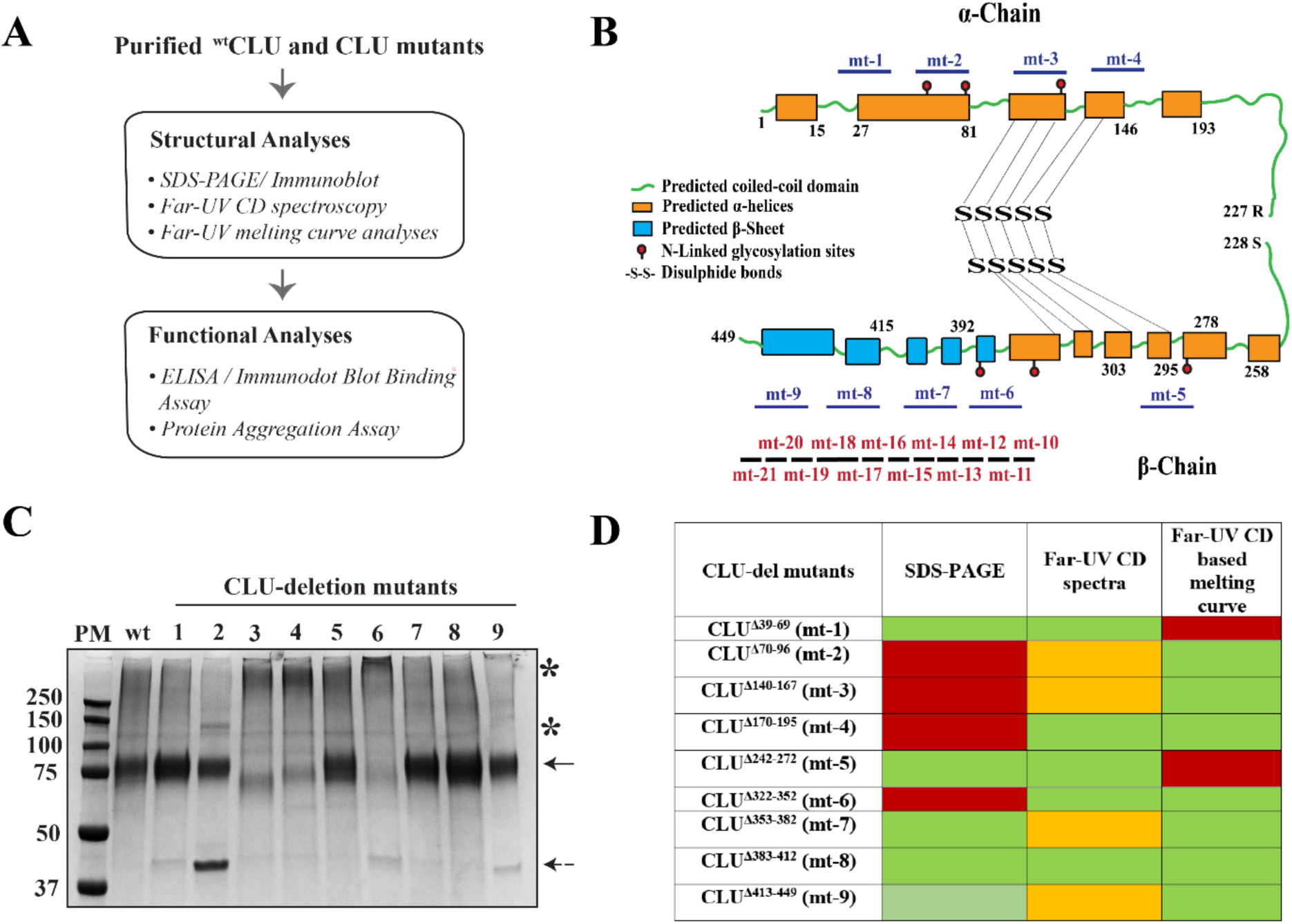
Summary of analyses of ^wt^CLU and CLU mutants. (A) Pipeline of structural and functional analyses used for characterisation of ^wt^CLU and CLU mutants. (B) Mature CLU has two chains (α and β) linked by five interchain disulphide bonds and is predicted to primarily composed of amphipathic α-helices followed by coiled-coil domains and beta sheets [20]. CLU also contains six glycosylation sites and R227 & S228 flank the proteolytic cleavage site of the precursor CLU molecule. Locations of residue deletions or alanine substitutions in the mature CLU molecule are indicated, respectively, for CLU-deletion mutants (blue; mt-1-mt-9) and alanine substitution mutants (red; mt-10-mt-21). (C) Non-reducing SDS-PAGE of ^wt^CLU and CLU-deletion mutants. Bands indicated with a solid arrow represent the expected mass of ^wt^CLU (∼ 75 kDa). Unexpected high molecular weight (HMW) or low-molecular weight (LMW) bands are indicated with asterisk or dashed arrow, respectively. The mass of individual bands (kDa) in a protein marker (PM) are indicated. Image is representative of 2 independent experiments. (D) Heat map summary of results for structural analyses comparing ^wt^CLU and CLU-deletion mutants. Colour scheme key: green (high similarity), orange (some differences) and red (substantial differences).

A total of nine deletion mutants were designed and expressed. The locations of the regions deleted (residues 39-69, 70-96, 140-167, 170-195, 242-272, 322-352, 352-382, 383-412 and 413-449; numbering includes the 22-mer signal peptide) were chosen to avoid cysteine residues involved in disulfide bonding, asparagine residues involved in N-linked glycosylation, and the inter-chain cleavage site. These nine mutants were designated mt-1 to mt-9, corresponding to the order of their positions proceeding from the N-terminus to the C-terminus of the CLU polypeptide (Figure 1B; Supplementary Table S1). All CLU mutants produced and recombinant ^wt^CLU were expressed in MEXi 293E cells (see Methods). Earlier studies aimed at comparing ^wt^CLU and CLU deletion mutants expressed the recombinant proteins bearing a twin Streptag positioned between residues 227 and 228 (as described in Supplementary Table S1) and purified the proteins using a Streptactin column as described [9]. In later studies working with alanine-stretch CLU mutants, these mutants and ^wt^CLU incorporated a C-terminal 4-residue C-tag (EPEA), and were purified by C-tag affinity chromatography [9].

### Structural analyses of CLU deletion mutants

#### SDS-PAGE analysis

Structural aberrations in CLU mutants frequently manifest as SDS-resistant high molecular weight (HMW) aggregates easily detected by SDS-PAGE. These HMW species may represent SDS/heat-resistant CLU oligomers that are each comprised of more than one α- and β-chain, possibly resulting from error(s) in the formation of inter-chain disulphide bonds. Purified CLU deletion mutants were analysed by non-reducing SDS PAGE. Being a heavily glycosylated glycoprotein, CLU migrates anomalously in SDS PAGE - it migrates with an apparent mass of 75-80 kDa although its actual mass is ∼ 60 kDa, and the major band resolved is broad owing to the presence of multiple glycovariants [6]. Similar to ^wt^CLU, mt -1, -5, -7, -8 and -9 showed a major band at ∼ 75 kDa (Figure 1C). Although the average mass of each individual deletion was ∼ 4000 Da (i.e. ∼ 7% of the total mass of CLU), the anomalous pattern of migration of CLU obscures these small differences in polypeptide mass. A minor band at ∼ 40 kDa was also detected for mt-2, and in lesser amounts for mt-1, 6 and 9 (Figure 1C). This band could represent non-disulfide-bonded single α or β chains, which previous studies have reported as resulting from the overexpression of ^wt^CLU in cancer cells [10]. A further minor band of unknown identity was also detected for mt-2 at ∼ 140 kDa. For mt-3, 4 and 6, the band detected at or near 75 kDa was significantly less intense and substantial HMW material was detected towards the top of the gel, consistent with aggregation (Figure 1C).

#### Far-UV CD analyses

Secondary structure content and melting points for ^wt^CLU and CLU deletion mutants were performed by far-UV circular dichroism (CD) spectroscopy. The far-UV CD spectra of mt-1, 4, 5 and 8 were similar to ^wt^CLU, however those of mt-2, 3, 6, 7 and 9 showed some differences (Supplementary Figure S1). In the latter group of mutants, the deletions span regions of predicted α-helix in the N-terminus of the α-chain and the C-terminus of the β-chain (Figure 1B), and thus may have impacted on their overall secondary structure content. Far-UV CD based melting curve analysis was next used to compare the unfolding patterns of ^wt^CLU and CLU deletion mutants. Mutant proteins with an overall decreased flexibility exhibit a delayed unfolding relative to the wild type counterpart [11]. The melting curves were broadly similar between ^wt^CLU, mt-2, mt-3, mt-6, mt-7, mt-8 and mt-9 (Supplementary Figure S2). Mt-1 and mt-5 appeared to be more resistant to melting than wt-CLU and did not substantively unfold even at 70 °C, while mt-4 showed changes in ellipticity even at temperatures below 40 °C (Supplementary Figure S2). Therefore, relative to ^wt^CLU, the thermal stability of mt-1, -4 and -5 are likely to be compromised.

On the basis of the combined results from both SDS-PAGE and Far-UV CD analyses, mutants mt-1 to mt-6 were identified as having substantial structural differences from ^wt^CLU and were therefore excluded from further analysis (Figure 1D). The remaining mutants (mt-7 to mt-9) were examined in functional analyses.

### Functional analyses of CLU deletion mutants

#### Protein aggregation assays

Previous studies have shown that CLU dose-dependently inhibits the in vitro amorphous aggregation of many different proteins including citrate synthase (CS), and the formation of amyloid by the 42-residue isoform of Aβ (Aβ^1–42^) [12]. The ability of ^wt^CLU and mt-7, mt-8 and mt-9 to inhibit the *in vitro* aggregation of CS and Aβ^1–42^ were compared in protein aggregation assays [13, 14]. Unlike ^wt^CLU, at a 1:6 molar ratio of CLU:CS, mt-7, mt-8, and mt-9 had very limited (or no) ability to inhibit CS aggregation (Figure 2A). Furthermore, relative to ^wt^CLU, at a 1:40 molar ratio of CLU:Aβ^1–42^, mt-7, mt-8 and mt-9 all showed a ∼ 50% reduced ability to inhibit endpoint Aβ^1–42^ aggregation (Figure 2B).

**Figure 2.**
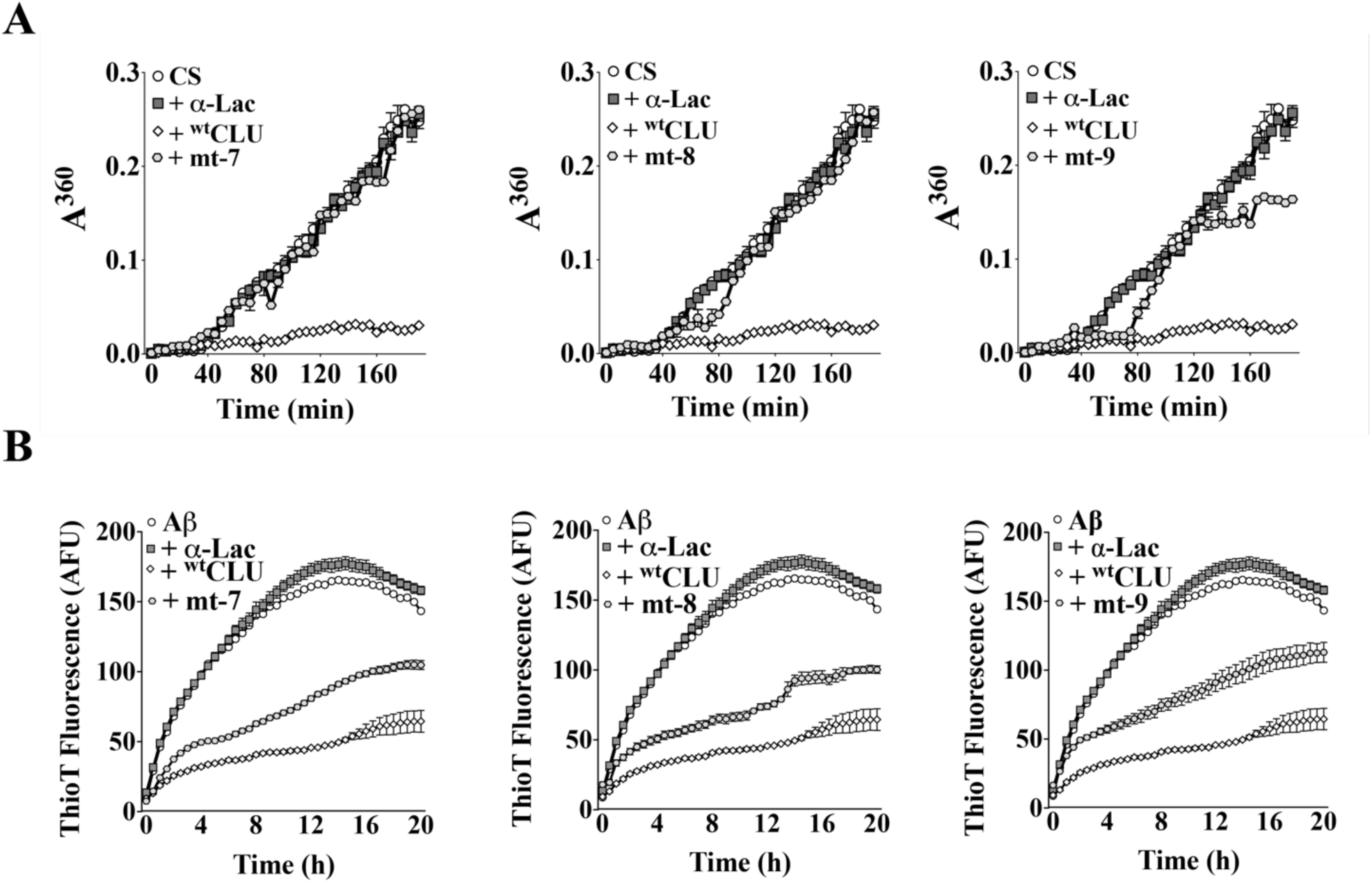
Comparison of the abilities of ^wt^CLU and CLU deletion mutants to inhibit the *in vitro* aggregation of CS and Aβ^1–42^. (A) Amorphous aggregation of CS was induced by incubation at 42 °C. CS (2 µM) was incubated with or without 0.3 µM of ^wt^CLU or mt-7, mt-8 or mt-9. 0.3 µM of α-Lactalbumin (α-Lac) was used as a non-chaperone control protein. Mean absorbances at 360 nm (A^360^) ± SEM (n = 4) are plotted. (B) Amyloid aggregation of Aβ^1–42^. 10 µM Aβ^1–42^ (Aβ), supplemented with 20 µM ThioT, was incubated with 0.25 µM of ^wt^CLU or mt-7, mt-8 or mt-9 for 20 h. 0.25 µM α-Lac was used as non-chaperone control protein. Mean ThioT fluorescence values (AFU, arbitrary fluorescence units) ± SEM (n = 4) are plotted. Error bars may be too small to be visible. In (A) and (B), α-Lac alone did not aggregate (not shown). Results shown are representative of 2 independent experiments.

#### CLU Ala-stretch mutants

The results of the analyses of CLU deletion mutants suggested that the C-terminal region of the CLU β-chain, which is spanned by the deletions corresponding to mt-7, mt-8 and mt-9 (Figure 1B), may contain binding regions that interact with misfolded client proteins. To investigate this possibility further, we designed and produced an additional series of twelve alanine-stretch (Ala-stretch) substitution mutants (mt-10 to mt-21) containing short stretches of 8-9 alanine residues spanning this region (Figure 1B, Supplementary Table S3). Alanine was selected for this purpose as it does not introduce any major steric or electrostatic effects that are likely to impact on overall protein conformation [15].

### Structural analyses of CLU Ala-stretch mutants

#### Immunoblot analyses

Given the potential for CLU mutations to cause structural changes observed in many of the deletion mutants, we first analysed culture supernatants harvested from transiently transfected MEXi293HEK cells expressing ^wt^CLU or Ala-stretch mutants by non-reducing immunoblot analysis. This procedure allowed us to quickly identify any CLU Ala-stretch mutants that formed incorrectly sized molecules. Reactive bands corresponding to the expected position of ^wt^CLU, were detected for mt-12, -13 and -21 (Supplementary Figure S3). However, for all other CLU Ala-stretch mutants, major bands were detected at ∼ 250 kDa or greater (Supplementary Figure S3), indicating aberrant aggregation. The variable band intensities detected are likely to result from differential expression of the various CLU mutants. Mutants mt-12, -13 and -21 were subsequently purified by C-tag affinity chromatography to yield different amounts of protein (Supplementary Figure S4 and Table S4) and subjected to further structural and functional characterisation.

#### Far-UV CD analyses

Far-UV CD spectroscopy was used to compare the secondary structure and thermal stability of ^wt^CLU and Ala-stretch mutants. Relative to ^wt^CLU, mt-12 and mt-21 had very similar far-UV CD spectra, while the spectrum between 205-220 nm (corresponding to alpha-helical content) and 220-230 nm (corresponding to beta-sheet content) for mt-13 was significantly lowered (Supplementary Figure S5A) [11]. The far-UV CD melting curves for mt-12, mt-13 and mt-21 were all very similar to that of ^wt^CLU (Supplementary Figure S5B). Collectively, the results of the far-UV CD analyses suggest that, relative to ^wt^CLU, the structures of mt-12 and mt-21 are very similar, and that mt-13 has a near-identical melting curve but lower α-helical and β-sheet content than ^wt^CLU. Despite its lowered α-helical and β-sheet content, on the basis of its striking similarity to ^wt^CLU in non-reducing SDS-PAGE and far-UV CD based melting curve analyses, we included mt-13, together with mt-12 and mt-21, in the functional analyses.

### Functional analyses of CLU Ala-stretch mutants

*In vitro* assays were performed to determine the relative abilities of mt-12, mt-13 and mt-21 versus ^wt^CLU to (i) bind to misfolded client proteins, and (ii) inhibit protein aggregation via either the amorphous or amyloid-forming pathways.

#### Binding to misfolded client proteins

##### Amorphously aggregating proteins

Previous studies have shown that CLU preferentially binds to misfolded glutathione-S-transferase (GST) and bovine serum albumin (BSA), both of which are known to aggregate amorphously [7]. Relative to ^wt^CLU, mt-12, mt-13 and mt-21 all showed significantly less binding to misfolded GST and BSA (Figure 3A, top panels). Relative to ^wt^CLU, the binding of mt-12 and mt-21 to either misfolded GST or BSA was reduced to a similar extent (by ∼ 75-90% and 90-95%, respectively), while that of mt-13 was reduced by ∼ 50% and ∼ 60%, respectively (Figure 3A, top panels). All CLU proteins showed very low levels of binding to both native GST and BSA (Figure 3A, bottom panels). *Amyloid-forming Aβ* ^1–42^. An immunodot blot assay was used to compare the binding of aggregated species of biotin-Aβ^1–42^ to CLU proteins (^wt^CLU, mt-12, mt-13, and mt-21) immobilised onto nitrocellulose membrane. Biotin-Aβ^1–42^ was collected from an *in vitro* aggregation assay mixture at 5 h (T1, pre-fibrillar, consisting mostly of Aβ^1-42^ oligomers) and 12 h (T2, fibrillar) after the start of aggregation [14] (Figure 3B, *top panel*). We previously reported that CLU binds preferentially to oligomeric Aβ [14]. As expected, negligible binding of fibrillar (T2) Aβ^1–42^ to any immobilized CLU protein was detected. In contrast, oligomeric Aβ^1–42^ (T1) bound strongly to ^wt^CLU but substantially less to mt-12, mt-13 and mt-21 (Figure 3B, *bottom panel*). Negligible binding of Aβ^1–42^ (T1 or T2) to the non-chaperone control protein α−lactalbumin was detected.

**Figure 3.**
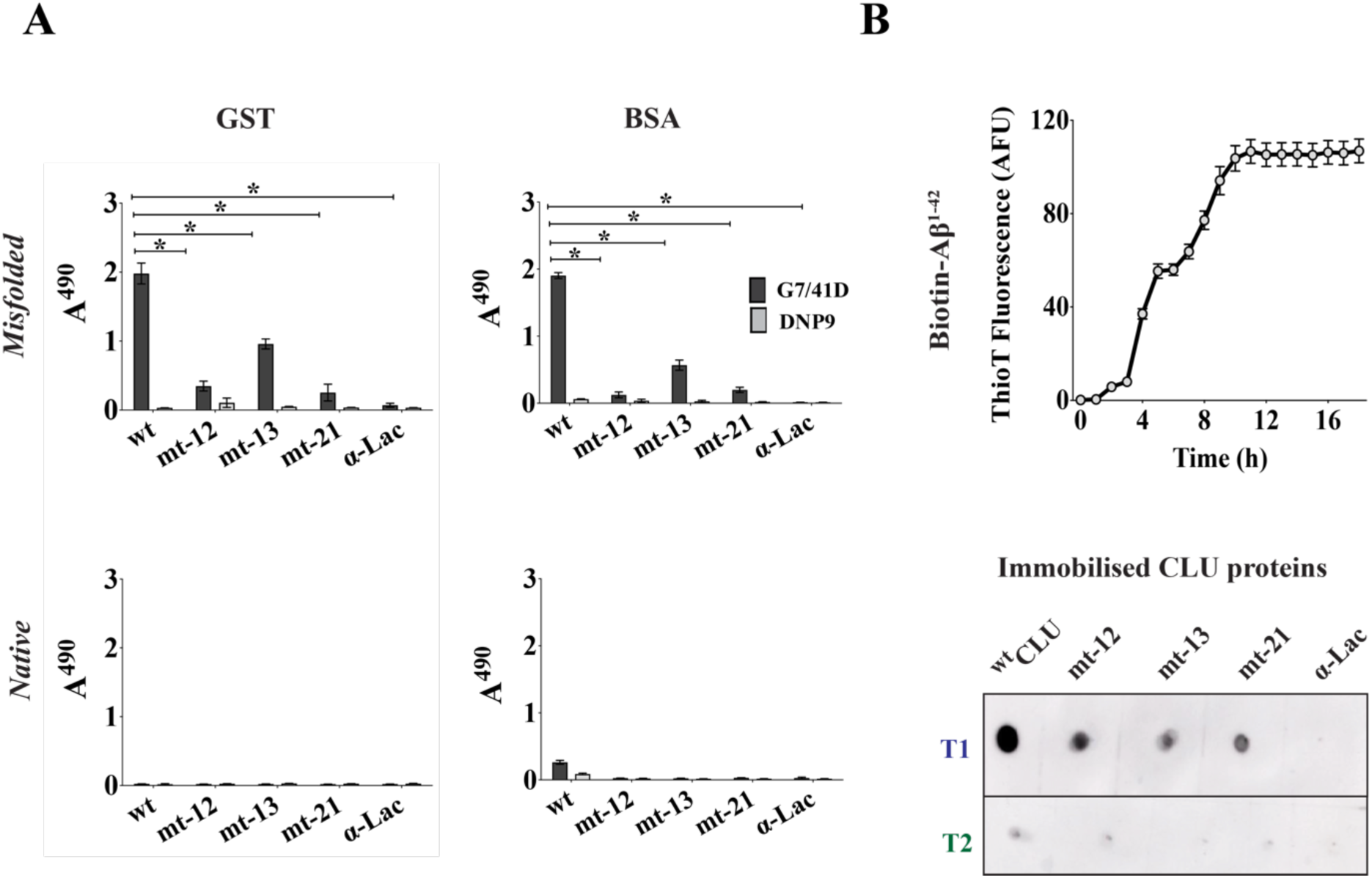
Ala-stretch CLU mutants show reduced interactions with chaperone client proteins. (A) ELISA measuring the binding of ^wt^CLU (wt) and mt-12, mt-13 & mt-21 to plate-immobilised GST (left panel) and BSA (right panel) client proteins. The client proteins were misfolded (upper panels) or native (lower panels). Bound CLU was detected using CLU specific antibody (1:1 mixture of G7 and 41D). An isotype matched control antibody (DNP9) was used as a control. Mean absorbances at 490 nm (A^490^) ± SEM (n = 4) are plotted. Asterisks indicate significant differences (Oneway ANOVA, p < 0.05). (B) *Upper panel*: Generation of pre-fibrillar and fibrillar Aβ^1–42^. Thioflavin T assay measuring the *in vitro* aggregation of biotin-Aβ^1–42^. Samples were collected at 5 h (T1; presumptive pre-fibrillar species) and 12 h (T2; presumptive fibrillar species) following the beginning of a biotin-Aβ^1–42^ aggregation assay. Lower panel: Immunodot blot assay measuring binding of CLU proteins to aggregated biotin-Aβ^1–42^ species. ^wt^CLU, mt-12, mt-13 or mt-21, and α-Lac (non-chaperone control protein) were immobilized on a nitrocellulose membrane and incubated with 10 µM biotin-Aβ^1–42^ (T1 or T2). Bound biotin-Aβ^1–42^ was detected using SA-HRP. In both (A) & (B) error bars are too small to be visible in some cases and results are representative of 2 independent experiments.

#### Protein aggregation assays

Relative to ^wt^CLU, mt-12 showed a ∼ 32% reduction in its ability to inhibit the end-point aggregation of CS, while the corresponding reductions for mt-13 and mt-21 were ∼ 92% and ∼ 68%, respectively (Figure 4A). The non-chaperone control protein α-lactalbumin did not significantly affect the endpoint level of CS aggregation (not shown). Similarly, relative to ^wt^CLU, the ability of mt-12 to inhibit the level of endpoint Aβ^1–42^ amyloid formation was reduced by ∼ 42%, and that of mt-13 and mt-21 both by ∼ 55% (Figure 4B). The non-chaperone control protein BSA did not significantly affect the endpoint level of Aβ^1–42^ amyloid formed (not shown).

**Figure 4.**
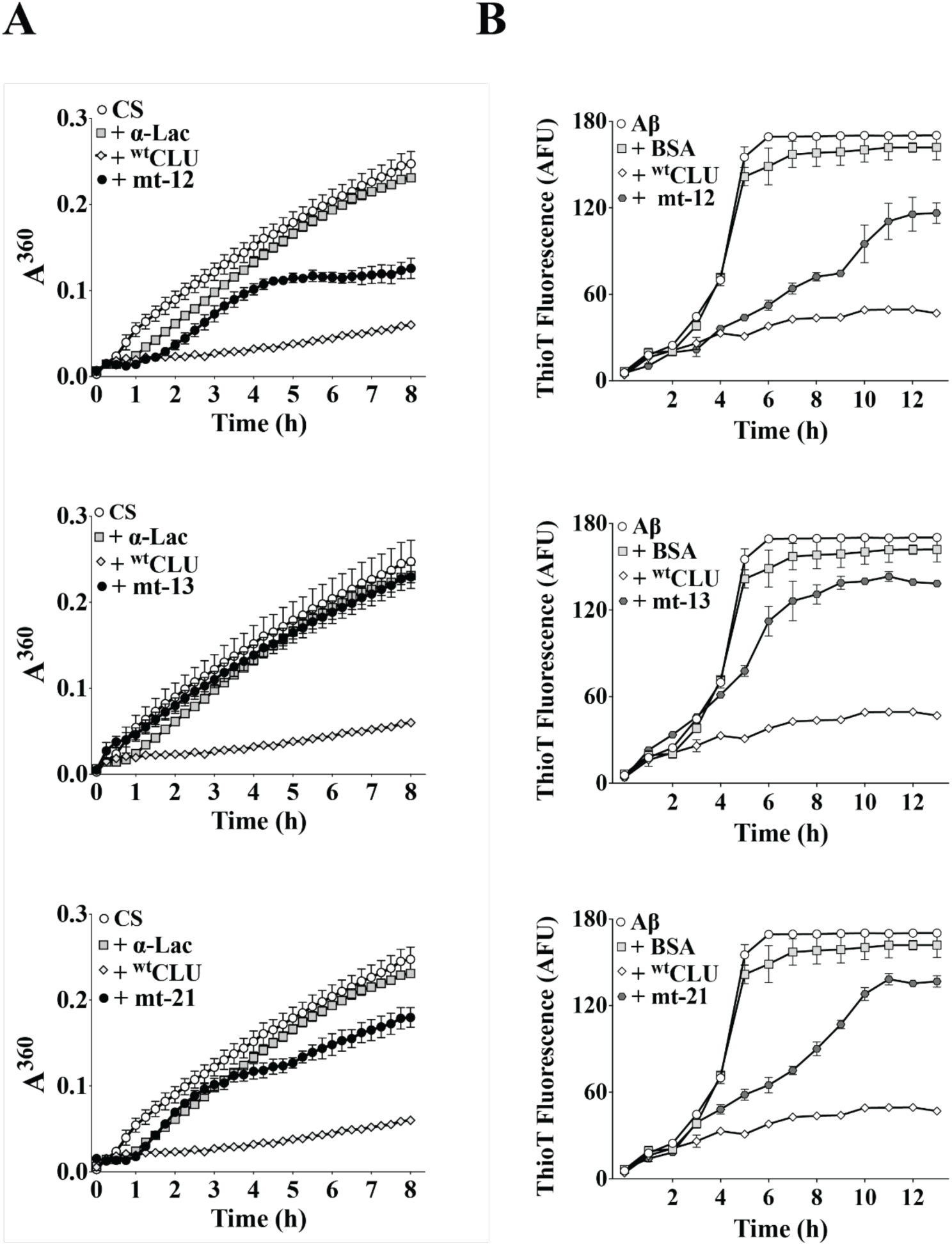
Comparing effects of ^wt^CLU and Ala-stretch mutants (mt-12, mt-13 & mt-21) on CS and Aβ^1–42^ aggregation. (A) Amorphous aggregation of CS. 2 µM CS was incubated with or without 1 µM ^wt^CLU, mt-12, mt-13, mt-21, or α-Lac (a non-chaperone control protein). When incubated alone under these conditions, α-Lac did not aggregate (not shown). Mean absorbances (AU) at 360 nm (A^360^) ± SEM (n = 4) are plotted. (B) *In vitro* Aβ^1–42^ aggregation *assay*. 10 µM Aβ^1–42^ (Aβ) was incubated with or without 0.25 µM ^wt^CLU, mt-12, mt-13, mt-21, or BSA (as a non-chaperone control protein). When incubated alone under these conditions, BSA did not aggregate (not shown). Mean ThioT fluorescence values (AFU, arbitrary fluorescence units x1000) ± SEM (n = 4) are plotted. In both (A) & (B) error bars are too small to be visible in some cases and results are representative of 2 independent experiments.

Collectively, the outcomes from the mutational analyses suggested that residues 369-384 (corresponding to mt-12 & mt-13) and 442-449 (corresponding to mt-21), positioned in the C-terminal region and at the C-terminus of the β-chain, respectively, may be important in the interactions of CLU with misfolded protein clients. However, it remained possible that the Ala-stretch mutations might induce changes in CLU structure outside the substituted regions that are not detected by the SDS PAGE and far-UV CD analyses, and that these effects are responsible for the observed inhibition of interactions with misfolded client proteins. To exclude this possibility, we used an independent approach to verify the involvement of these two β-chain regions in the chaperone activity of CLU – *in silico* designed camelid nanobodies were used as probes for the function of these regions.

### Sequence-specific designed nanobodies verify CLU β-chain regions as client interaction sites

Owing to their small size (∼ 14 kDa) and flexible extended CD3 binding loop, the binding of *in silico* designed sequence-specific nanobodies (dNbs) to targeted epitopes on a protein of interest introduces minimal steric interference [16]. Consequently, dNbs can be used as powerful tools to decipher protein-protein interactions. Using a previously published algorithm for the prediction of putative binding regions for dNbs, *in silico* analysis of the CLU sequence suggested a high probability of success in designing dNb binding sequences specific for CLU residues 369-376 and 442-449 (corresponding to mt-12 and mt-21, respectively), but not for CLU residues 377-384 (corresponding to mt-13) [16]. Consequently, we expressed and purified dNbs specific for CLU residues 369-376 and 442-449 (dNb^369–376^ and dNb^442–449^), together with another dNb containing a scrambled binding sequence not expected to bind to CLU (dNb^neg^; amino acid sequence details for all dNbs listed in Supplementary Table S5). Like the recombinant ^wt^CLU and mutant CLU molecules, all dNb incorporated a C-terminal C-tag to permit single-step affinity purification.

The specificity of the dNbs were validated in immuno-dot blot assays in which the binding of the dNbs to human plasma derived CLU immobilized onto nitrocellulose membrane was specifically inhibited by a molar excess of peptides corresponding to the designed target sequence, but not control peptides of different sequence (data not shown). We next tested if the dNbs targeting CLU residues 369-376 and 442-449 could specifically reduce the ability of CLU to inhibit the amorphous aggregation of CS and Aβ^1–42^ amyloid formation *in vitro*. Pre-incubation of ^wt^CLU with dNb^369–376^ or dNb^442–449^ reduced its ability to inhibit the end point levels of CS aggregation by ∼ 96 % and ∼ 70 %, respectively (Figure 5A) and that of Aβ^1–42^ amyloid formation by ∼ 60 % and ∼ 100 %, respectively (Figure 5B). In contrast, pre-incubation of ^wt^CLU with dNb^neg^ had no significant effect on its ability to inhibit the *in vitro* aggregation of either CS or Aβ^1–42^ (Figures 5A & 5B). Furthermore, pre-incubation with dNb^369–376^ or dNb^442–449^ did not affect the *in vitro* aggregation of either CS or Aβ^1–42^ (not shown). Lastly, the non-chaperone control protein BSA did not affect the aggregation of CS or Aβ^1–42^ (Figures 5A & 5B) and did not aggregate itself during the experiment (not shown).

**Figure 5.**
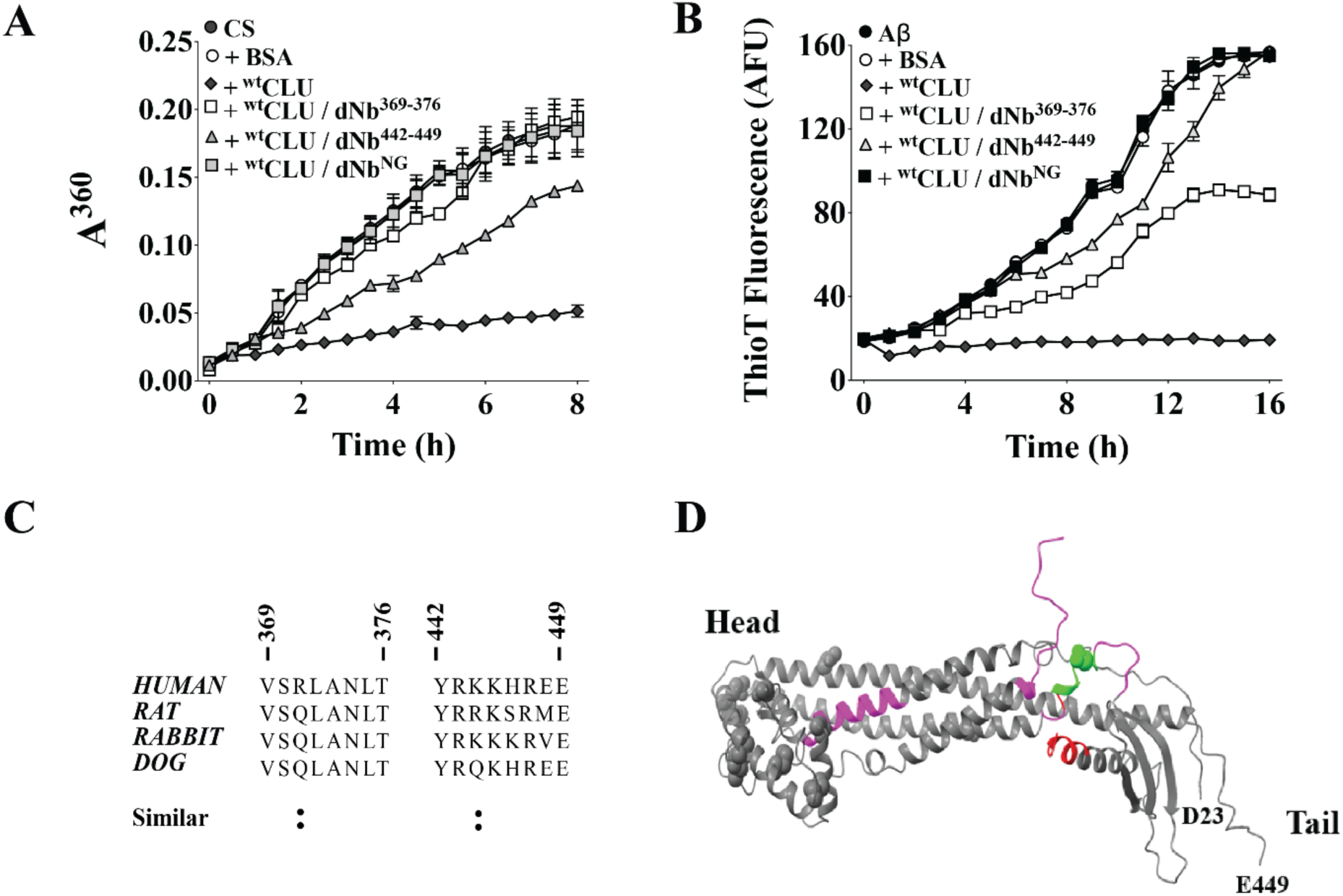
Effects of sequence-specific dNbs on the ability of CLU to inhibit aggregation of CS and Aβ^1–42^, binding site sequence conservation, and *in silico* predicted binding sites on CLU. (A and B) Relative to untreated ^wt^CLU, ^wt^CLU pre-incubated with dNb^369–376^ or dNb^442–449^ (but not dNb^neg^) had a reduced ability to inhibit the aggregation of (A) CS or (B) Aβ^1–42^ amyloid aggregation (measured using thioflavin T (ThT) fluorescence). In (A) 2 µM CS was incubated alone or with (i) 1 µM ^wt^CLU, (ii) 1 µM ^wt^CLU pre-incubated with 1 µM dNb^369–376^ or dNb^442–449^ or dNb^NG^, or (iii) 1 µM dNb^369–376^ or dNb^442–449^ or dNb^NG^. Mean absorbances (AU) at 360 nm ± SEM (n = 4) are plotted. In (B) 10 µM Aβ^1–42^ (Aβ) was incubated alone or with (i) 0.25 µM ^wt^CLU, (ii) 0.25 µM ^wt^CLU pre-incubated with 0.25 µM dNb^369–376^ or dNb^442–449^, or or dNb^NG^ (iii) 0.25 µM dNb^369–376^ or dNb^442–449^ or dNb^NG^. Mean ThioT fluorescence values (AFU, arbitrary fluorescence units x1000) ± SEM (n = 4) are plotted. BSA (non-chaperone control protein) or dNb^369–376^ or dNb^442–449^ used at 1 µM in (A) and 0.25 µM in (B), aggregated when incubated alone (not shown). Some error bars are too small to be visible and results shown are representative of 2 independent experiments. (C) CLU residues 369-376 (corresponding to mt-12) and 442-449 (corresponding to mt-21) are highly conserved between species, being identical between human, rat, rabbit and dog with the exception of a single chemically similar residue Q/R/K at positions 371 and 444 (indicated by colons). Sequences were obtained from UniProtKB and sequence conservation was assessed using Clustal Omega multiple sequence alignment [21]. (D) Predicted and experimentally validated CLU ligand binding sites. The 3D structure of CLU (in grey) predicted using AlpaFold (PDB ID: AF-P10909-F1) was used for *in silico* protein-protein interaction studies to identify cryptic binding sites using FTMove [17]. D23 (aspartate at position 23) and E449 (glutamate at position 449) indicate the N- and C-termini of the mature CLU molecule; disulphide-bonded head and flexible tail region indicated. Color-coded regions indicate (i) purple – predicted ligand binding sites (ii) red - experimentally verified ligand binding sites, and (iii) lime green-regions are both predicted and experimentally verified as ligand binding sites.

The results of the sequence-specific dNb studies confirm the suggestion from the mutational analyses that CLU residues 369-376 and 442-449 are important in its interactions with misfolded client proteins and its ability to inhibit their aggregation *in vitro*.

## Discussion

In the absence of an experimentally-determined structure for CLU, but guided by the AlphaFold predicted structure (Figure 5D), we initially applied mutagenesis to “scan” parts of the CLU molecule to identify regions likely to be important in its interactions with chaperone client proteins. We used a recently developed mammalian expression system to produce ^wt^CLU which is structurally and functionally very similar to plasma-derived CLU [9]. The same system was used to produce a variety of CLU mutants. A CLU mutant could possess a compromised chaperone activity either as a result of structural aberrations, or from the deletion/disruption of a client protein binding site. Therefore, we aimed to identify CLU mutants that were structurally similar to ^wt^CLU but which had compromised chaperone activity.

We initially deleted parts of the CLU molecule and subjected the mutants to structural assays to identify any with aberrant structures, which were excluded from subsequent functional assays. The design of the nine CLU deletion mutants was relatively conservative, each comprising ∼ 7% of the total mass of the protein, and positioned to avoid interference with inter-chain disulfide bonding, N-linked glycosylation of asparagine residues, and the inter-chain cleavage site (Supplementary Table S2). Despite this strategy, when compared with ^wt^CLU, 6 of these 9 mutants showed structural aberrations. These results suggest that the fidelity of the complex post-translational processing of CLU has quite limited tolerance for modifications to the native sequence, and highlights the crucial importance of verifying the structure of recombinant CLU mutants before subjecting them to functional analyses.

The results of SDS PAGE and far UV CD analyses identified CLU deletion mutants mt-7 to mt-9 as being structurally similar to ^wt^CLU (Figures 1C & 1D, Supplementary Figures S1 & S2). Therefore, these three mutants (all bearing deletions in the C-terminal region of the CLU β-chain) were subsequently analysed in functional assays. Mt-7 to mt-9 showed a complete or almost complete loss of their ability to inhibit the amorphous aggregation of CS, and ∼ 50% loss of their ability to inhibit the aggregation of Aβ^1–42^ to form amyloid (Figure 2A & 2B). These results suggested that the C-terminal region of the CLU β-chain (Figure 1B) is likely to contain potential binding sites for chaperone client proteins.

To test this hypothesis, we next generated twelve CLU Ala-stretch mutants in which stretches of 8-9 alanine residues were substituted along the CLU molecule between residues 352-449 (Supplementary Table S3). Immunoblot analyses of culture supernatants containing overexpressed CLU Ala-stretch mutants showed that all except three (mt-12, mt-13 & mt-21) formed HMW species, suggesting structural aberrations (Supplementary Figure S3). Additional structural characterisation indicated that, compared to ^wt^CLU, all three mutants had similar thermal stability, and mt-12 and mt-21 (not mt-13) also shared similar secondary structure content (Supplementary Figure S4). Subsequently functional assays were performed to interrogate whether Ala-substitution in the regions of CLU sequence targeted by mt-12, mt-13 and mt-21 (CLU residues 369-376, 377-384, and 442-449, respectively) contain partial or complete binding sites important for its interaction with misfolded client proteins.

When compared with ^wt^CLU, all three β-chain Ala-substitution mutants tested showed significant reductions in their abilities to bind to (i) misfolded GST and BSA, and (ii) biotin-Aβ^1–42^. Furthermore, these mutants also showed reduced abilities to inhibit the endpoint aggregation of (i) CS (reductions of ∼ 32% for mt-12, ∼ 92% for mt-13, ∼ 68% for mt-21) and Aβ^1–42^ (reductions of ∼ 42% for mt-12, and ∼ 55% for mt-13 & mt-21) (Figures 3 & 4).

Collectively, the mutational analyses suggested that residues 369-384 (positioned in the C-terminal tail of the β-chain, corresponding to mt-12 & mt-13) and 442-449 (representing the β-chain C-terminus, corresponding to mt-21) are likely to represent partial (or complete) binding sites for misfolded protein clients. This observation was confirmed with the use of sequence-specific dNbs. *In silico* analysis of the CLU sequence suggested the likelihood that designing dNbs specific for CLU residues 369-376 and 442-449 (corresponding to mt-12 and mt-21, respectively) would be successful, but that targeting CLU residues 377-384 (corresponding to mt-13) would be more difficult. For this reason, we generated dNbs targeting CLU residues 369-376 (dNb^369–376^) and 442-449 (dNb^442–449^) and showed that when pre-incubated with ^wt^CLU these dNb (but not the control dNb^neg^) specifically and substantially reduced the ability of CLU to inhibit the *in vitro* aggregation of CS and Aβ^1–42^. The reductions in the ability of CLU to inhibit CS and Aβ^1–42^ endpoint aggregation were, respectively, ∼ 100% and ∼ 64% for dNb^369–376^, and ∼ 74% and ∼ 100% for dNb^442–449^ (Figures 5A&B). Therefore, both mutational analyses and blocking experiments performed with sequence-specific dNb strongly implicate CLU residues 369-376 and 442-449 as being key binding sites important for its interactions with misfolded protein client molecules. These two stretches of CLU residues are 85-100% conserved between different mammalian species (Figure 5C), consistent with these regions being important in the biological functions of CLU.

The limited tolerance of CLU to mutational changes meant that we were forced to exclude many mutants from functional analyses on the basis of detectable structural changes. Thus, it is very possible that CLU contains additional misfolded protein client binding sites distinct from the two identified in the current study, and positioned in other parts of the CLU molecule. The observation that Ala-substitution of 369-376 or 442-449 resulted in substantial but not complete loss of chaperone activity (Figures 3B & 4B) supports this hypothesis. Even in the cases when, under the conditions tested, specific dNbs binding almost abolished the ability of CLU to inhibit endpoint protein aggregation, there was nevertheless some evidence that CLU still partly slowed the kinetics of aggregation (Figures 5A&B). Therefore, it is likely that the two misfolded client protein binding sites reported here are critically involved in the interactions of CLU with misfolded proteins, but their action may be complemented by other yet to be experimentally verified binding sites located elsewhere in the molecule.

Lastly, we used FTMove [17], a machine-learning algorithm to predict functional binding sites on CLU and compare those to the sites that were experimentally validated here. FTMove was trained with validated binding sites of ∼ 7622 proteins from the Protein Data Bank database, enabling high-confidence simulation and prediction of unknown binding sites on protein of interest using a distance and volumetric overlap approach. FTMove analyses predicted four CLU ligand binding sites, including S370-Y382 and N363-W368 which overlap or are in close proximity to our identified binding region at residues 369-376 (Figure 5D, Supplementary Table S6). The remaining FTMove predicted ligand binding sites S228-A242 and N157-L172 (Figure 5D, Supplementary Table S6) are distinct from those identified in the current study, suggesting that these sites may bind to other CLU client proteins or to cell surface receptors.

Collectively, the locations of the experimentally identified and *in silico*-predicted ligand binding sites are consistent with a model in which the binding of multiple flexible regions in the CLU tail (Figure 5D) enables it to interact with structurally diverse misfolding protein clients. In contrast, the predicted ligand binding site N157-L162 is located distally from the flexible tails, positioned in proximity to the disulfide-bonded head, where it may function as a cell receptor binding site. Physical separation of client protein and cell receptor binding sites could act to minimize any potential steric hindrance of the latter when CLU-client protein complexes bind to cells prior to their uptake and degradation. However, the locations of CLU binding sites for cell surface receptors remain to be confirmed.

In conclusion, we anticipate that the growing understanding of the relationship between CLU structure and function may enable the future development of rationally designed, engineered CLU with an increased ability to neutralize and safely dispose of specific toxic aggregating extracellular proteins. These forms of CLU could provide unique new therapeutics for the many serious human diseases arising from the accumulation of toxic and pro-inflammatory extracellular protein aggregates.

## Materials and Methods

### Materials

α-Lactalbumin (α-Lac), bovine serum albumin (BSA), glutathione-S transferase (GST), citrate synthase (CS), thimerosal, iodoacetamide, casein, hydrogen peroxide (H_2_O_2_), citric acid, thioflavin-T (Thio-T), Greiner 6-well cell culture plates, Erlenmeyer non-baffled cell culture shaker flasks, Greiner 96-well clear flat bottom ELISA plates, and Greiner 384-well clear flat bottom plates were purchased from Sigma Aldrich (Australia). Aβ^1–42^ was purchased from ANASPEC Australia and biotin-Aβ^1–42^ was purchased from Sapphire Biosciences (Australia). All other chemicals were of analytical grade and purchased from Sigma Aldrich (Australia) unless otherwise specified.

### Bacterial culture

*Escherichia coli* (*E. coli)* strains were used for plasmid propagation and extraction. Chemically competent DH5α (*E. coli*) cells (purchased from NEB, Australia) were used for transformation with plasmid DNA as per the manufacturer’s protocol, plated on LB agar plates containing 100 μg/ml ampicillin after transformation and grown overnight at 37 °C. Single bacterial colonies picked from these plates were inoculated into liquid LB culture containing 100 μg/ml ampicillin and grown with shaking (180-200 rpm) at 37 ° C for 16-18 h. Finally, the bacterial culture was harvested by centrifugation at 10000 x g for 10 min at 4 °C for subsequent extraction of plasmid DNA using the CompactPrep plasmid kit, following the manufacturer’s protocol (Qiagen, Australia). Expression plasmids encoding dNb were transformed into BL21 T7 Lys *E. coli* cells and periplasmic expression was induced with 100 μM IPTG, as described previously [18].

### Plasmid designs

#### CLU mutants

Insert DNA sequences coding for ^wt^CLU, nine deletion mutants (with a twin-Streptag (ST) positioned at the C-terminus of the α-chain), or twelve alanine-stretch mutants (with a C-tag^120^ (CT) positioned at the β-chain C-terminus) were produced by gene synthesis prior to cloning into pcDNA3.1 (Gene Universal, USA). Sequence details of all the plasmids used in this study are shown in Supplementary Tables S1 and S3. The deletions were 33-37 residues in length and each was positioned a minimum of 6 residues distant from regions involved in post-translational modifications (e.g. signal peptide cleavage, the inter-chain cleavage site, and the disulfide-bonded head region) (Figure 1B; Supplementary Table S1). Each deletion mutant retained at least 5 of the 6 N-linked glycosylation sites. It was shown previously that enzymatic removal of all glycosylation from CLU does not impair its chaperone activity [19]. Alanine-stretch mutants contained alanine substitutions of 8-9 residues in length positioned in the C-terminal region of the CLU β-chain (residues 352-449). Each Ala-stretch was at least 30 residues distant from regions involved in post-translational modifications, and each Ala-stretch mutant contained all 6 endogenous N-linked glycosylation sites (Figure 1B; Supplementary Table S3). DNA sequences for the dNbs including the CDR3 extended loop, binding region and a C-tag (for affinity purification) were produced by gene synthesis prior to cloning into the pET28(b)+ vector (Gene Universal, USA). Sequence details for dNb expression plasmids used in this study are shown in Supplementary Table S5.

### Recombinant protein expression and purification

^wt^CLU and all CLU mutants were expressed in suspension-cultured MEXi 293E cells [9]. Routine culture and transfection of MEXi 293E cells were performed following the manufacturer’s instructions (IBA Life Sciences, Germany). ***Strep-tagged ^wt^CLU and CLU deletion mutants***. Culture supernatants were first dialysed x3 against phosphate buffered saline (PBS; 137 mM NaCl, 2.7 mM KCl, 7.9 mM Na_2_HPO_4_, 1.5 mM KH_2_PO_4_, pH 7.4) and then supplemented with 1/10^th^ volume of 10X buffer W (1 M Tris, 1.5 M NaCl, 10 mM EDTA, pH 8.0) and 1x protease inhibitor cocktail tablets (as per the manufacturer’s instructions) before loading 200-600 ml sample onto a 5 ml bed volume Streptactin-XT affinity column (IBA Life Sciences, Germany). Purified protein was eluted using buffer BXT (1X buffer W containing 0.05 M biotin). ***C-tagged Ala-stretch CLU mutants and dNb*.** Culture supernatants were supplemented with 20 mM Tris buffer and 1x protease inhibitor cocktail tablets, adjusted to pH 8.0, and then loaded onto a 5 ml bed volume Capture Select C-tag XL affinity column (Thermo Fisher Scientific, Australia). Purified protein was eluted with 20 mM Tris, 2 M MgCl_2_, pH 8.0, dialysed x3 against PBS and then concentrated using Amicon Ultra15 centrifugal filters (Merck Millipore, Australia) with a molecular weight cut-off of 15 kDa (CLU) or 3.5 kDa (dNb). The absorbance at 280 nm (A^280^) of purified proteins was measured using a SPECTROstar Nano plate reader (BMG Labtech, Australia) and the concentrations calculated using extinction coefficients of 0.84 (0.1% A^280^) for CLU and *in-silico* calculated values provided in Supplementary Table S5 for dNb. All purified proteins were stored at -20°C.

### SDS PAGE

Protein samples were supplemented with an equal volume of 2X SDS sample buffer (120 mM Tris, 2% (w/v) SDS, 20% (v/v) glycerol, 0.02% (w/v) bromophenol blue, pH 6.8) and heated for 5 min at 95 °C. Samples were then loaded onto 4-12% Bolt Bis-Tris gels (Thermo Fisher Scientific, Australia). Electrophoresis was conducted at 150-200 V for 30 min at room temperature (RT) in a Mini Gel tank using MES-SDS running buffer (Thermo Fisher Scientific, Australia). All gels were stained using InstantBlue Coomassie protein stain (Expedon, Australia) and imaged with a ChemiDoc XRS+ System (Bio-Rad, Australia).

### Immunoblotting

Samples were electrophoretically transferred from gels onto nitrocellulose membranes (Pall Corporation, Pensacola, FL, USA) overnight at 30 V and 4 °C using Western transfer buffer (25 mM Tris-Cl, 192 mM glycine, 20% v/v ethanol, pH 7.6) in a Trans Blot apparatus (Bio-Rad, Australia). Blots were then blocked for 1 h at room temperature using 5% (w/v) skimmed milk powder in PBS (SM/PBS). C-tagged proteins were detected using Capture biotin anti-C tag conjugate (Thermo Fisher Scientific, Australia) followed by Streptavidin-HRP (Abcam, Australia), both used at a 1:2000 dilution in SM/PBS overnight at 4 °C. Between detection reagent incubations, blots were washed x3 with PBS/0.1% (v/v) Triton X-100 followed by 2 washes with PBS. Blots were then developed with a pico chemiluminescence (ECL) kit (Thermo Fisher Scientific, Australia) and imaged using an Amersham 600 RGB imager (GE Healthcare, Australia). In immunodot blot assays, small volumes of protein samples (2-12 µl) were spotted onto nitrocellulose membrane and processed as described for immunoblotting.

### Far-UV circular dichroism (CD) spectroscopy

Far-UV CD spectroscopy of ^wt^CLU and CLU mutants was performed using a Jasco Model J-810 spectropolarimeter connected to a CDF-426S/L Peltier system (Jasco, Australia). Secondary structure content was assessed by measuring the 190-240 nm CD spectra of purified protein samples at 0.1 mg/ml in 5 mM phosphate buffer, pH 7.4, in a 0.1 cm path length cuvette (Sigma Aldrich, Australia). Results were collected as an average of 6 scans at 37 °C with a bandwidth and data pitch of 1 nm, high sensitivity (5 mdeg), and using a continuous scanning mode at 100 nm/min. For the melting curve analyses, CD spectra at 222 nm were measured using a 10-70 °C ramp (in increments of 1 degree) and the results were acquired as an average of 3 scans with data pitch of 0.5 nm, bandwidth of 1 nm, and high sensitivity (5 mdeg). Buffer only (5 mM phosphate, pH 7.4) was used to correct the raw values for the contribution of the buffer.

### ELISA

A previously established ELISA method was used to detect the binding of ^wt^CLU and CLU mutants to native or misfolded GST and BSA. GST (20 μg/ml in PBS, 50 µl/ well) was adsorbed onto the wells of two 96 well ELISA plates for 1 h at 37 °C. One of these plates was then heated at 60 °C for 1 h (to induce misfolding of GST) and the other plate was stored at 4°C (native GST plate) until further use. BSA (1 mg/ml in PBS, 50 µl/well) was adsorbed onto the wells of two 96 well ELISA plates either supplemented with (misfolded BSA plate) or without (native BSA plate) 20 mM DTT and incubated overnight at 4 °C. Subsequently, to block any potentially reactive free sulfhydryl groups, both plates were incubated for 1 h at 37°C with 100 µl/well of 5 mM iodoacetamide in PBS. After washing, all the plates were then blocked for 1 h at RT with 1% (w/v) heat denatured casein (HDC), 0.01% (w/v) thimerosal in PBS (HDC/PBS). Subsequently, 13 μg/ml of purified ^wt^CLU or mutant CLU diluted in HDC/PBS was added to each well (50 µl/well) and incubated for 1 h at RT. Bound CLU was detected with a 1:1 mixture of G7 and 41D hybridoma culture supernatants (containing monoclonal antibodies specific for human CLU), and DNP9 mouse hybridoma culture supernatant was used as a source of isotype matched control antibody [7]. Bound primary antibodies were detected by incubation with 50 µl/well of goat anti-mouse IgG HRP (Agilent Technologies, Inc., USA) diluted 1:2000 in HDC/PBS. All antibody incubations were for 1 h at room temperature and the ELISA plates were washed x6 with MilliQ water between incubations. Finally, 50 µl of substrate (2.5 mg/ml o-phenylenediamine dihydrochloride in 0.05 M citric acid, 0.1 M Na_2_HPO_4_, 0.03% (v/v) H_2_O_2_, pH 5.0) was added to each well and the reaction stopped when required using 100 µl/well of 1M hydrochloric acid (HCl). Absorbance at 490 nm was measured using a SPECTROstar Nano Plate reader (BMG Labtech, Australia).

### *In vitro* protein aggregation assays

#### CS aggregation assay

CS (2 μM, 50 μl/well) in 50 mM Tris, 8 mM HEPES, pH 8.0 was heated for 4-5 h at 43 °C in a Greiner 384 flat bottom clear-well plate. ^wt^CLU, CLU mutants, or α-Lac (non-chaperone control protein) were added to a final concentration of 0.25 or 0.5 μM. Each sample was prepared in triplicate and the absorbance at 360 nm (A^360^) measured in a SPECTROstar Nano plate reader (BMG Labtech, Australia) with 20 light flashes per reading and 3 s double orbital shaking between each read (1 mm shaking width, 600 rpm). In these assays, the same volume of buffer only was used to correct the raw absorbance values for each sample. *Aβ*^1–42^ *aggregation assay.* Aβ^1–42^ (10 µM in PBS, 50 µl/well, supplemented with 20 µM Thio-T) was incubated in a Greiner 384 flat bottom plate for 14-16 h at 30 °C without shaking. ^wt^CLU, CLU mutants, or α-Lac (non-chaperone control protein) were added to a final concentration of 0.25 or 0.5 μM. Each sample was prepared in triplicate and the Thio-T fluorescence emission measured in a POLARstar plate reader (BMG Labtech, Australia) equipped with 440/10 nm (excitation) and 520/25 nm (emission) band pass filters. In these assays, a matching volume of PBS only was used to correct the raw fluorescence values for each sample.

### Statistical Analysis

Statistical analyses of data were performed using Oneway-ANOVA with Bonferonni comparison between each pair of samples and P values < 0.05 were considered to be statistically significant.

### Author Contributions

S.S. performed most of the experimental work and contributed to authoring of the manuscript. Y.S. expressed and purified recombinant CLU proteins. E.P. expressed, purified and tested dNb. P.S. and M.V. performed *in silico* designs of dNb and edited the manuscript. M.R.W. coordinated and supervised all experimental work and edited the manuscript.

## Acknowledgements

SS is grateful to the Australian government for an Australian Postgraduate Award. YS thanks the China Scholarship Council for their support.

