## Supplementary Materials for "Two sites in the C-terminal β-chain tail mediate interactions of the chaperone clusterin with amyloid beta and other misfolded client proteins"

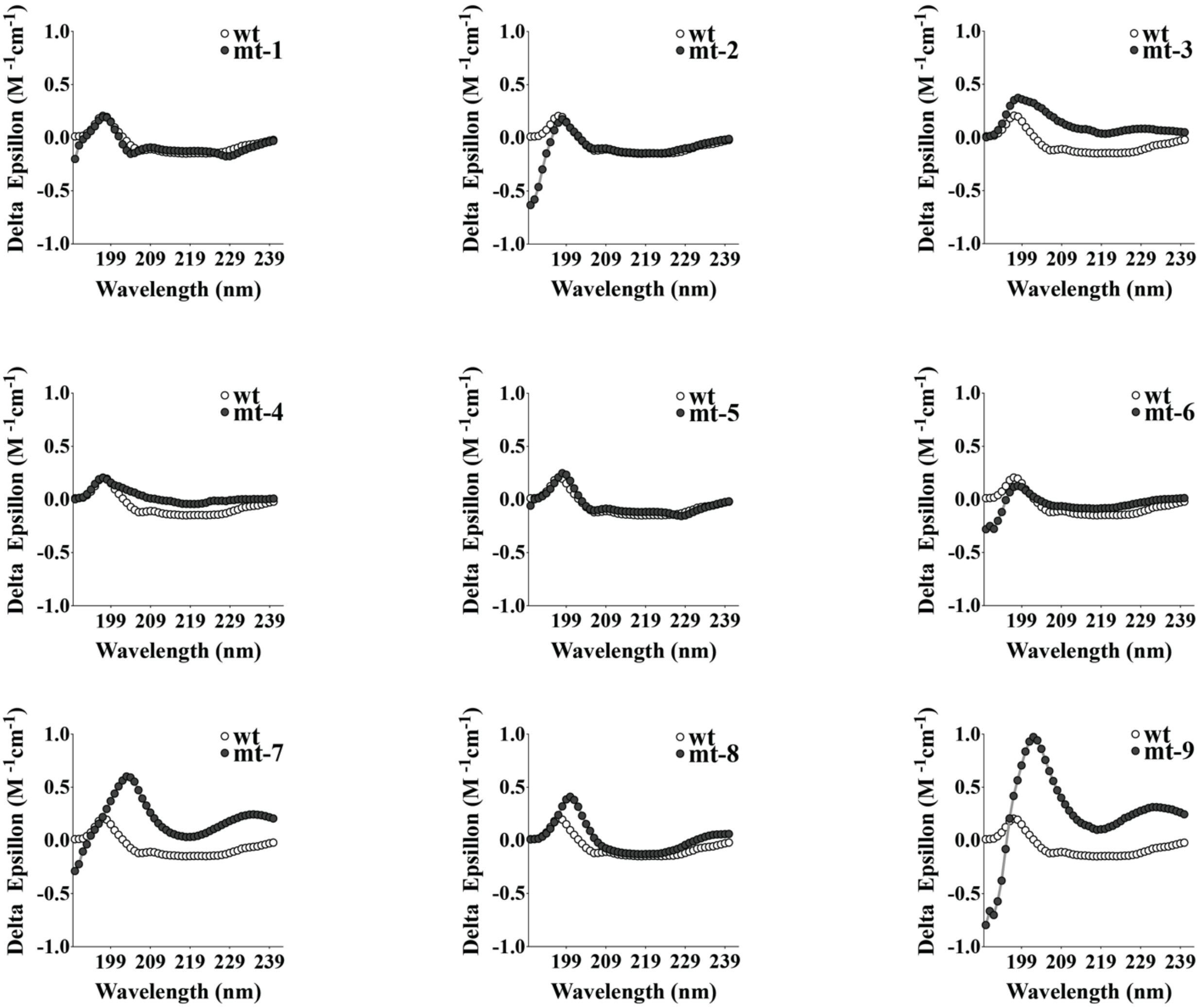


**Supplementary Figure S1. Far-UV CD spectra of $^{wt}$CLU (wt) and CLU deletion mutants.** Purified $^{wt}$CLU and mt-1 to mt-9 were analysed between 190-240 nm. Means ± SEM (n =10) are plotted, error bars are too small to be visible. Results shown here represent a single experiment.

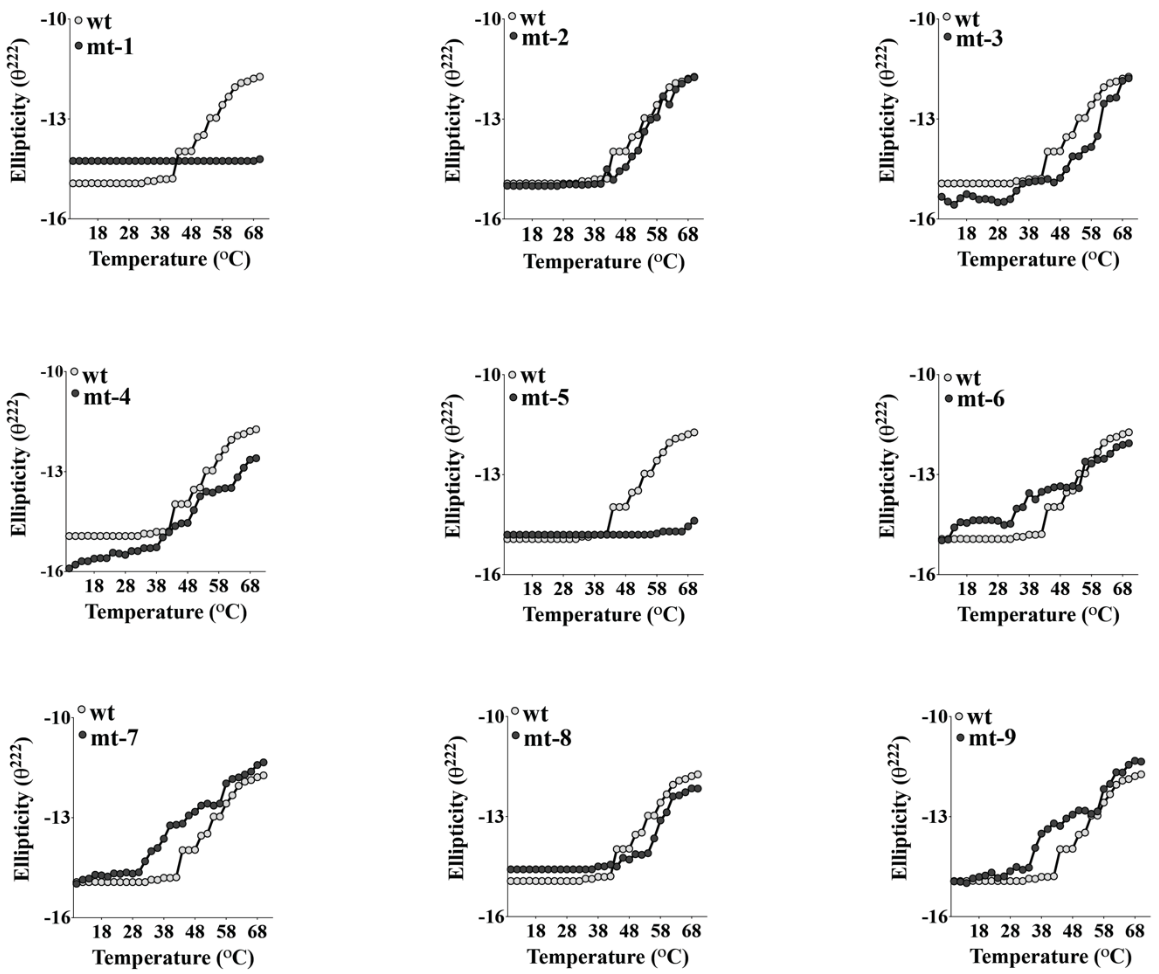


**Supplementary Figure S2. Far-UV CD based melting curve analyses of $^{wt}$CLU (wt) and CLU-deletion mutants.** Purified proteins were analysed at 222 nm between 10-70 °C. Means ± SEM (n = 3) are plotted, error bars are too small to be visible. Results shown here are representative of a single experiment.

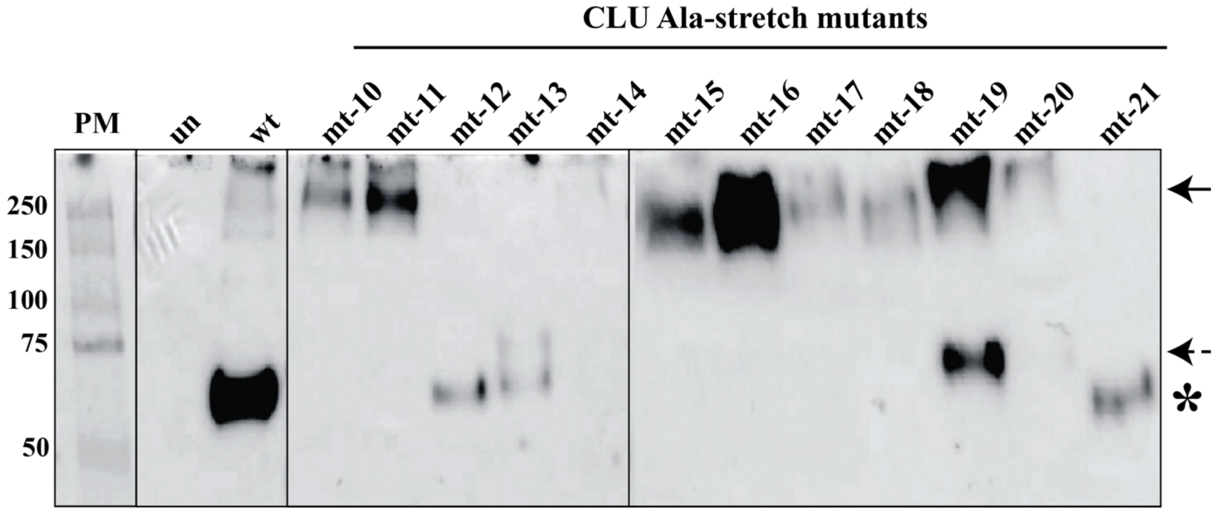


**Supplementary Figure S3. Immunoblot analysis of culture supernatants containing wtCLU and Ala-stretch mutants.** Culture supernatants were harvested from untransfected (un) or transfected cells overexpressing wtCLU (wt) or Ala-stretch mutants (mt-10 to mt-21). Proteins were separated by non-reducing SDS-PAGE prior to electrophoretic transfer to a membrane and the blot probed with biotin anti-C tag antibody. The masses of individual bands (kDa) in a protein marker (PM, *left lane*) are indicated. The asterisk indicates the expected mass of wtCLU under these conditions. Solid arrow indicates high molecular weight (HMW) complexes at ~230-270 kDa; dashed arrow indicates a species at ~75 kDa; asterisk indicates the position of wtCLU at ~ 70 kDa. Image shown here is representative of 2 independent experiments.

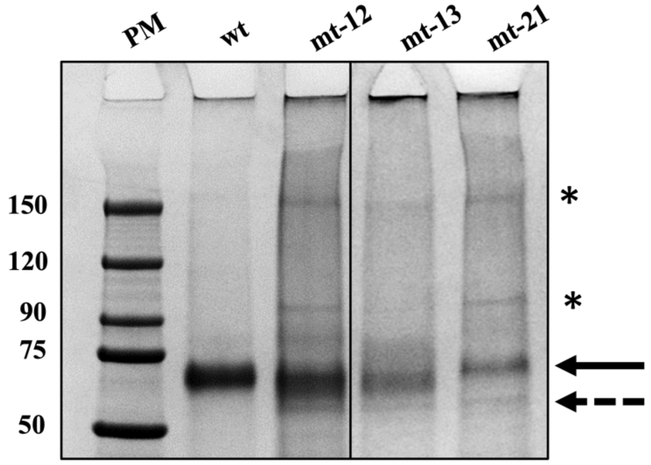


**Supplementary Figure S4. Non-reducing SDS-PAGE of $^{wt}$CLU (wt) and Ala-stretch mutants.** Bands indicated with a solid arrow represent the expected mass of $^{wt}$CLU (wt). Unexpected high molecular weight (HMW) or low-molecular weight (LMW) bands are indicated with an asterisk or dashed arrow, respectively. The masses of individual bands (kDa) in a protein marker (PM) are indicated. Image is representative of 2 independent experiments.

**A**

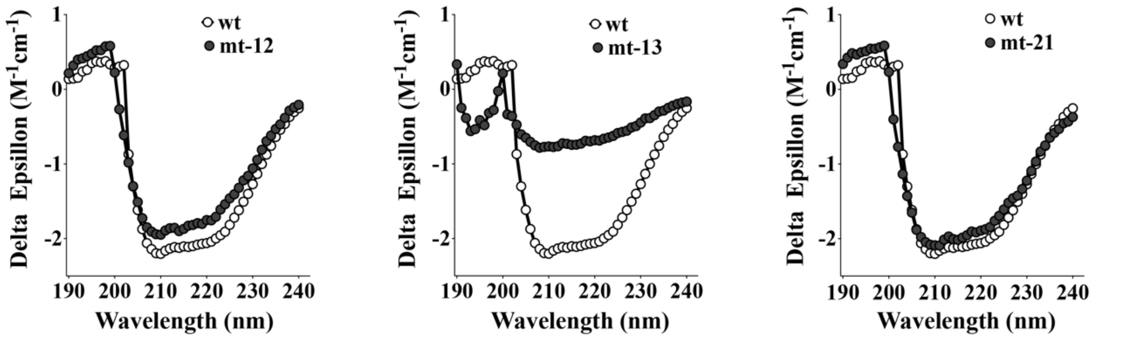


**B**

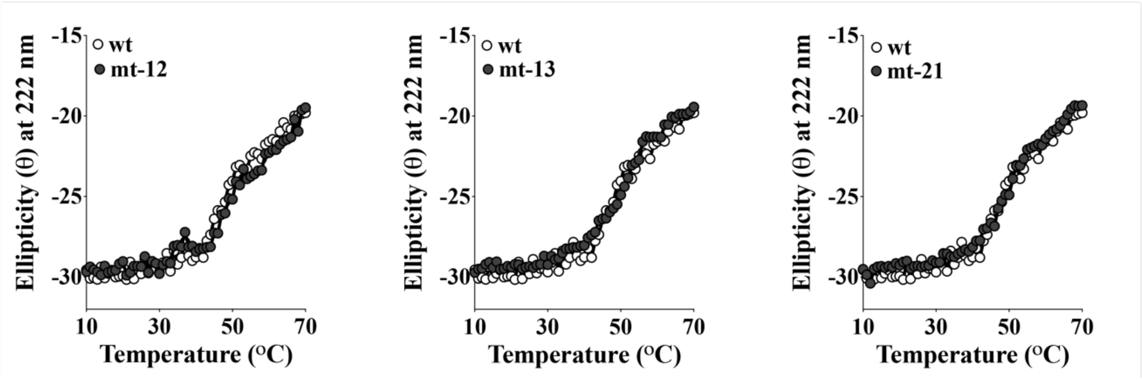


**Supplementary Figure S5. Far-UV CD analyses of purified $^{wt}$CLU (wt) and Ala-stretch mutants. (A) Far-UV CD spectra recorded between 190-240 nm are plotted as** mean delta epsilon (molar extinction coefficient) ± SEM (n = 10). (B) Far-UV CD based melting curve analysis of proteins measured between 10-70 °C, plotted as mean ellipticity values at 222 nm ± SEM (n = 3). In both cases error bars are too small to be visible. Results shown are from a single experiment.

**Table S1 Amino acid sequences of CLU proteins encoded by pCDNA3.1 (+) plasmids.**

| Plasmid Name | Sequence |
|---|---|
| $^{wt}$CLU-ST | MMKTLLLFVGLLLTWESGQVLGDQTVSDNELQEMSNQGSKYV<br>NKEIQNAVNGVKQIKTLIEKTNEERKTLLSNLEEAKKKKEDALN<br>ETRESETKLKELPGVCNETMMALWEECKPCLKQTCMKFYARVC<br>RSGSGLVGRQLEEFLNQSSPFYFWMNGDRIDSLLENDRQQTHM<br>LDVMQDHFSRASSIIDELFQDRFFTREPQDTYHYLPFSLPHRRPH<br>FFFPKSGGGGSA**WSHPQFEK*****GGGSGGGSGGSA*****WSHPQFEK**GG<br>GGRIVRSLMPFSPYEPLNFHAMFQPFLEMIHEAQQAMDIHFHSPA<br>FQHPPTEFIREGDDDRTVCREIRHNSTGCLRMKDQCDKCREILSV<br>DCSTNNPSQAKLRRELDESLQVAERLTRKYNELLKSYQWKMLN<br>TSSLLEQLNEQFNWVSRLANLTQGEDQYYLRVTTVASHTSDSDV<br>PSGVTEVVVKLFDSDPITVTVPVEVSRKNPKFMETVAEKALQEY<br>RKKHREE |
| CLU$^{\Delta 39\text{-}69}$-ST<br>(mt-1) | MMKTLLLFVGLLLTWESGQVLGDQTVSDNELQEMSNQGLLSNL<br>EEAKKKKEDALNETRESETKLKELPGVCNETMMALWEECKPCL<br>KQTCMKFYARVCRSGSGLVGRQLEEFLNQSSPFYFWMNGDRIDS<br>LLENDRQQTHMLDVMQDHFSRASSIIDELFQDRFFTREPQDTYH<br>YLPFSLPHRRPHFFFPKSGGGGSA**WSHPQFEK*****GGGSGGGSGGS***<br>***A*****WSHPQFEK**GGGGRIVRSLMPFSPYEPLNFHAMFQPFLEMIHEA<br>QQAMDIHFHSPAFQHPPTEFIREGDDDRTVCREIRHNSTGCLRM<br>KDQCDKCREILSVDCSTNNPSQAKLRRELDESLQVAERLTRKYN<br>ELLKSYQWKMLNTSSLLEQLNEQFNWVSRLANLTQGEDQYYLR<br>VTTVASHTSDSDVPSGVTEVVVKLFDSDPITVTVPVEVSRKNPKF<br>METVAEKALQEYRKKHREE |
| CLU$^{\Delta 70\text{-}96}$-ST<br>(mt-2) | MMKTLLLFVGLLLTWESGQVLGDQTVSDNELQEMSNQGSKYV<br>NKEIQNAVNGVKQIKTLIEKTNEERKTELPGVCNETMMALWEEC<br>KPCLKQTCMKFYARVCRSGSGLVGRQLEEFLNQSSPFYFWMNG<br>DRIDSLLENDRQQTHMLDVMQDHFSRASSIIDELFQDRFFTREPQ<br>DTYHYLPFSLPHRRPHFFFPKSGGGGSA**WSHPQFEK*****GGGSGGG***<br>***SGGSA*****WSHPQFEK**GGGGRIVRSLMPFSPYEPLNFHAMFQPFLE<br>MIHEAQQAMDIHFHSPAFQHPPTEFIREGDDDRTVCREIRHNSTG<br>CLRMKDQCDKCREILSVDCSTNNPSQAKLRRELDESLQVAERLT<br>RKYNELLKSYQWKMLNTSSLLEQLNEQFNWVSRLANLTQGED<br>QYYLRVTTVASHTSDSDVPSGVTEVVVKLFDSDPITVTVPVEVS<br>RKNPKFMETVAEKALQEYRKKHREE |
| CLU$^{\Delta 140\text{-}167}$-ST<br>(mt-3) | MMKTLLLFVGLLLTWESGQVLGDQTVSDNELQEMSNQGSKYV<br>NKEIQNAVNGVKQIKTLIEKTNEERKTLLSNLEEAKKKKEDALN<br>ETRESETKLKELPGVCNETMMALWEECKPCLKQTCMKFYARVC<br>RSGSGLVGRQQQTHMLDVMQDHFSRASSIIDELFQDRFFTREPQ<br>DTYHYLPFSLPHRRPHFFFPKSGGGGSA**WSHPQFEK*****GGGSGGG*** |

| | |
|---|---|
| | *SGGSA***WSHPQFEK**GGGGRIVRSLMPFSPYEPLNFHAMFQPFLE<br>MIHEAQQAMDIHFHSPAFQHPPTEFIREGDDDRTVCREIRHNSTG<br>CLRMKDQCDKCREILSVDCSTNNPSQAKLRRELDESLQVAERLT<br>RKYNELLKSYQWKMLNTSSLLEQLNEQFNWVSRLANLTQGED<br>QYYLRVTTVASHTSDSDVPSGVTEVVVKLFDSDPITVTVPVEVS<br>RKNPKFMETVAEKALQEYRKKHREE |
| CLU$^{\Delta 170-195}$-ST (mt-4) | MMKTLLLFVGLLLTWESGQVLGDQTVSDNELQEMSNQGSKYV<br>NKEIQNAVNGVKQIKTLIEKTNEERKTLLSNLEEAKKKKEDALN<br>ETRESETKLKELPGVCNETMMALWEECKPCLKQTCMKFYARVC<br>RSGSGLVGRQLEEFLNQSSPFYFWMNGDRIDSLLENDRQQFTRE<br>PQDTYHYLPFSLPHRRPHFFFPKSGGGGSA**WSHPQFEK***GGGSG*<br>*GGSGGSA***WSHPQFEK**GGGGRIVRSLMPFSPYEPLNFHAMFQPF<br>LEMIHEAQQAMDIHFHSPAFQHPPTEFIREGDDDRTVCREIRHNS<br>TGCLRMKDQCDKCREILSVDCSTNNPSQAKLRRELDESLQVAER<br>LTRKYNELLKSYQWKMLNTSSLLEQLNEQFNWVSRLANLTQGE<br>DQYYLRVTTVASHTSDSDVPSGVTEVVVKLFDSDPITVTVPVEV<br>SRKNPKFMETVAEKALQEYRKKHREE |
| CLU$^{\Delta 242-272}$-ST (mt-5) | MMKTLLLFVGLLLTWESGQVLGDQTVSDNELQEMSNQGSKYV<br>NKEIQNAVNGVKQIKTLIEKTNEERKTLLSNLEEAKKKKEDALN<br>ETRESETKLKELPGVCNETMMALWEECKPCLKQTCMKFYARVC<br>RSGSGLVGRQLEEFLNQSSPFYFWMNGDRIDSLLENDRQQTHM<br>LDVMQDHFSRASSIIDELFQDRFFTREPQDTYHYLPFSLPHRRPH<br>FFFPKSGGGGSA**WSHPQFEK***GGGSGGGSGGSA***WSHPQFEK**GG<br>GGRIVRSLMPFSPYEPLNFHEFIREGDDDRTVCREIRHNSTGCLR<br>MKDQCDKCREILSVDCSTNNPSQAKLRRELDESLQVAERLTRKY<br>NELLKSYQWKMLNTSSLLEQLNEQFNWVSRLANLTQGEDQYY<br>LRVTTVASHTSDSDVPSGVTEVVVKLFDSDPITVTVPVEVSRKNP<br>KFMETVAEKALQEYRKKHREE |
| CLU$^{\Delta 322-352}$-ST (mt-6) | MMKTLLLFVGLLLTWESGQVLGDQTVSDNELQEMSNQGSKYV<br>NKEIQNAVNGVKQIKTLIEKTNEERKTLLSNLEEAKKKKEDALN<br>ETRESETKLKELPGVCNETMMALWEECKPCLKQTCMKFYARVC<br>RSGSGLVGRQLEEFLNQSSPFYFWMNGDRIDSLLENDRQQTHM<br>LDVMQDHFSRASSIIDELFQDRFFTREPQDTYHYLPFSLPHRRPH<br>FFFPKSGGGGSA**WSHPQFEK***GGGSGGGSGGSA***WSHPQFEK**GG<br>GGRIVRSLMPFSPYEPLNFHAMFQPFLEMIHEAQQAMDIHFHSPA<br>FQHPPTEFIREGDDDRTVCREIRHNSTGCLRMKDQCDKCREILSV<br>DCSTNNPSQALNTSSLLEQLNEQFNWVSRLANLTQGEDQYYLR<br>VTTVASHTSDSDVPSGVTEVVVKLFDSDPITVTVPVEVSRKNPKF<br>METVAEKALQEYRKKHREE |
| | MMKTLLLFVGLLLTWESGQVLGDQTVSDNELQEMSNQGSKYV<br>NKEIQNAVNGVKQIKTLIEKTNEERKTLLSNLEEAKKKKEDALN<br>ETRESETKLKELPGVCNETMMALWEECKPCLKQTCMKFYARVC |

| | |
|---|---|
| CLU$^{\Delta 352\text{-}382}$-ST (mt-7) | RSGSGLVGRQLEEFLNQSSPFYFWMNGDRIDSLLENDRQQTHM LDVMQDHFSRASSIIDELFQDRFFTREPQDTYHYLPFSLPHRRPH FFFPKSGGGGSA**WSHPQFEK***GGGSGGGSGGSA***WSHPQFEK**GG GGRIVRSLMPFSPYEPLNFHAMFQPFLEMIHEAQQAMDIHFHSPA FQHPPTEFIREGDDDRTVCREIRHNSTGCLRMKDQCDKCREILSV DCSTNNPSQAKLRRELDESLQVAERLTRKYNELLKSYQWKYLR VTTVASHTSDSDVPSGVTEVVVKLFDSDPITVTVPVEVSRKNPKF METVAEKALQEYRKKHREE |
| CLU$^{\Delta 383\text{-}412}$-ST (mt-8) | MMKTLLLFVGLLLTWESGQVLGDQTVSDNELQEMSNQGSKYV NKEIQNAVNGVKQIKTLIEKTNEERKTLLSNLEEAKKKKEDALN ETRESETKLKELPGVCNETMMALWEECKPCLKQTCMKFYARVC RSGSGLVGRQLEEFLNQSSPFYFWMNGDRIDSLLENDRQQTHM LDVMQDHFSRASSIIDELFQDRFFTREPQDTYHYLPFSLPHRRPH FFFPKSGGGGSA**WSHPQFEK***GGGSGGGSGGSA***WSHPQFEK**GG GGRIVRSLMPFSPYEPLNFHAMFQPFLEMIHEAQQAMDIHFHSPA FQHPPTEFIREGDDDRTVCREIRHNSTGCLRMKDQCDKCREILSV DCSTNNPSQAKLRRELDESLQVAERLTRKYNELLKSYQWKMLN TSSLLEQLNEQFNWVSRLANLTQGEDQYDPITVTVPVEVSRKNP KFMETVAEKALQEYRKKHREE |
| CLU$^{\Delta 413\text{-}449}$-ST (mt-9) | MMKTLLLFVGLLLTWESGQVLGDQTVSDNELQEMSNQGSKYV NKEIQNAVNGVKQIKTLIEKTNEERKTLLSNLEEAKKKKEDALN ETRESETKLKELPGVCNETMMALWEECKPCLKQTCMKFYARVC RSGSGLVGRQLEEFLNQSSPFYFWMNGDRIDSLLENDRQQTHM LDVMQDHFSRASSIIDELFQDRFFTREPQDTYHYLPFSLPHRRPH FFFPKSGGGGSA**WSHPQFEK***GGGSGGGSGGSA***WSHPQFEK**GG GGRIVRSLMPFSPYEPLNFHAMFQPFLEMIHEAQQAMDIHFHSPA FQHPPTEFIREGDDDRTVCREIRHNSTGCLRMKDQCDKCREILSV DCSTNNPSQAKLRRELDESLQVAERLTRKYNELLKSYQWKMLN TSSLLEQLNEQFNWVSRLANLTQGEDQYYLRVTTVASHTSDSDV PSGVTEVVVKLFDS |

**Notes:**

1. Key:
   MMKTLLLFVGLLLTWESGQVLG: Signal peptide sequence
   GGGG: 4 glycine spacer sequence
   SA: Serine alanine spacer sequence
   **WSHPQFEK**: Single Streptag sequence
   ***GGGSGGGSGGSA:*** Internal spacer sequence between two Streptag sequences
2. The superscript (e.g. $^{\Delta 413\text{-}449}$) indicates those residues in mature CLU that have been deleted. In all these plasmids, twin-Streptag (ST) was positioned at the C-terminus of the CLU α-chain.
3. Identification codes for different CLU deletion proteins (mt-1-mt-9) are shown in parentheses under the full name.

**Table S2 Yield of purified CLU-ST proteins transiently expressed in MEXi 293E cells.**

| Plasmid Name | Yield (mg/L) |
|---|---|
| $^{wt}$CLU-ST | 25.0 |
| CLU$^{\Delta 39\text{-}69}$-ST (mt-1) | 2.8 |
| CLU$^{\Delta 70\text{-}96}$-ST (mt-2) | 0.3 |
| CLU$^{\Delta 140\text{-}167}$-ST (mt-3) | 15.4 |
| CLU$^{\Delta 170\text{-}195}$-ST (mt-4) | 3.8 |
| CLU$^{\Delta 242\text{-}272}$-ST (mt-5) | 10.0 |
| CLU$^{\Delta 322\text{-}352}$-ST (mt-6) | 1.1 |
| CLU$^{\Delta 352\text{-}382}$-ST (mt-7) | 2.9 |
| CLU$^{\Delta 383\text{-}412}$-ST (mt-8) | 20.0 |
| CLU$^{\Delta 413\text{-}449}$-ST (mt-9) | 0.8 |

**Table S3 Amino acid sequences of $^{wt}$CLU and alanine-stretch CLU mutants.**

| Plasmid Name | Sequence |
|---|---|
| $^{wt}$CLU-CT | MMKTLLLFVGLLLTWESGQVLGDQTVSDNELQEMSNQGSK<br>YVNKEIQNAVNGVKQIKTLIEKTNEERKTLLSNLEEAKKKKE<br>DALNETRESETKLKELPGVCNETMMALWEECKPCLKQTCMK<br>FYARVCRSGSGLVGRQLEEFLNQSSPFYFWMNGDRIDSLLEN<br>DRQQTHMLDVMQDHFSRASSIIDELFQDRFFTREPQDTYHYL<br>PFSLPHRRPHFFFPKSRIVRSLMPFSPYEPLNFHAMFQPFLEMI<br>HEAQQAMDIHFHSPAFQHPPTEFIREGDDDRTVCREIRHNSTG<br>CLRMKDQCDKCREILSVDCSTNNPSQAKLRRELDESLQVAER<br>LTRKYNELLKSYQWKMLNTSSLLEQLNEQFNWVSRLANLTQ<br>GEDQYYLRVTTVASHTSDSDVPSGVTEVVVKLFDSDPITVTVP<br>VEVSRKNPKFMETVAEKALQEYRKKHREE**EPEA** |
| CLU-Ala$^{352-360}$-CT (mt-10) | MMKTLLLFVGLLLTWESGQVLGDQTVSDNELQEMSNQGSK<br>YVNKEIQNAVNGVKQIKTLIEKTNEERKTLLSNLEEAKKKKE<br>DALNETRESETKLKELPGVCNETMMALWEECKPCLKQTCMK<br>FYARVCRSGSGLVGRQLEEFLNQSSPFYFWMNGDRIDSLLEN<br>DRQQTHMLDVMQDHFSRASSIIDELFQDRFFTREPQDTYHYL<br>PFSLPHRRPHFFFPKSRIVRSLMPFSPYEPLNFHAMFQPFLEMI<br>HEAQQAMDIHFHSPAFQHPPTEFIREGDDDRTVCREIRHNSTG<br>CLRMKDQCDKCREILSVDCSTNNPSQAKLRRELDESLQVAER<br>LTRKYNELLKSYQWKM**AAAAAAAA**QLNEQFNWVSRLANLT<br>QGEDQYYLRVTTVASHTSDSDVPSGVTEVVVKLFDSDPITVT<br>VPVEVSRKNPKFMETVAEKALQEYRKKHREE**EPEA** |
| CLU-Ala$^{361-368}$-CT (mt-11) | MMKTLLLFVGLLLTWESGQVLGDQTVSDNELQEMSNQGSK<br>YVNKEIQNAVNGVKQIKTLIEKTNEERKTLLSNLEEAKKKKE<br>DALNETRESETKLKELPGVCNETMMALWEECKPCLKQTCMK<br>FYARVCRSGSGLVGRQLEEFLNQSSPFYFWMNGDRIDSLLEN<br>DRQQTHMLDVMQDHFSRASSIIDELFQDRFFTREPQDTYHYL<br>PFSLPHRRPHFFFPKSRIVRSLMPFSPYEPLNFHAMFQPFLEMI<br>HEAQQAMDIHFHSPAFQHPPTEFIREGDDDRTVCREIRHNSTG<br>CLRMKDQCDKCREILSVDCSTNNPSQAKLRRELDESLQVAER<br>LTRKYNELLKSYQWKMLNTSSLLE**AAAAAAAA**VSRLANLTQ<br>GEDQYYLRVTTVASHTSDSDVPSGVTEVVVKLFDSDPITVTVP<br>VEVSRKNPKFMETVAEKALQEYRKKHREE**EPEA** |
| CLU-Ala$^{369-376}$-CT (mt-12) | MMKTLLLFVGLLLTWESGQVLGDQTVSDNELQEMSNQGSK<br>YVNKEIQNAVNGVKQIKTLIEKTNEERKTLLSNLEEAKKKKE<br>DALNETRESETKLKELPGVCNETMMALWEECKPCLKQTCMK<br>FYARVCRSGSGLVGRQLEEFLNQSSPFYFWMNGDRIDSLLEN<br>DRQQTHMLDVMQDHFSRASSIIDELFQDRFFTREPQDTYHYL<br>PFSLPHRRPHFFFPKSRIVRSLMPFSPYEPLNFHAMFQPFLEMI |

| | |
|---|---|
| | HEAQQAMDIHFHSPAFQHPPTEFIREGDDDRTVCREIRHNSTG<br>CLRMKDQCDKCREILSVDCSTNNPSQAKLRRELDESLQVAER<br>LTRKYNELLKSYQWKMLNTSSLLEQLNEQFNWAAAAAAAA<br>QGEDQYYLRVTTVASHTSDSDVPSGVTEVVVKLFDSDPITVT<br>VPVEVSRKNPKFMETVAEKALQEYRKKHREE**EPEA** |
| CLU-Ala$^{377-384}$-CT<br>(mt-13) | MMKTLLLFVGLLLTWESGQVLGDQTVSDNELQEMSNQGSK<br>YVNKEIQNAVNGVKQIKTLIEKTNEERKTLLSNLEEAKKKKE<br>DALNETRESETKLKELPGVCNETMMALWEECKPCLKQTCMK<br>FYARVCRSGSGLVGRQLEEFLNQSSPFYFWMNGDRIDSLLEN<br>DRQQTHMLDVMQDHFSRASSIIDELFQDRFFTREPQDTYHYL<br>PFSLPHRRPHFFFPKSRIVRSLMPFSPYEPLNFHAMFQPFLEMI<br>HEAQQAMDIHFHSPAFQHPPTEFIREGDDDRTVCREIRHNSTG<br>CLRMKDQCDKCREILSVDCSTNNPSQAKLRRELDESLQVAER<br>LTRKYNELLKSYQWKMLNTSSLLEQLNEQFNWVSRLANLTA<br>AAAAAAARVTTVASHTSDSDVPSGVTEVVVKLFDSDPITVTV<br>PVEVSRKNPKFMETVAEKALQEYRKKHREE**EPEA** |
| CLU-Ala$^{385-392}$-CT<br>(mt-14) | MMKTLLLFVGLLLTWESGQVLGDQTVSDNELQEMSNQGSK<br>YVNKEIQNAVNGVKQIKTLIEKTNEERKTLLSNLEEAKKKKE<br>DALNETRESETKLKELPGVCNETMMALWEECKPCLKQTCMK<br>FYARVCRSGSGLVGRQLEEFLNQSSPFYFWMNGDRIDSLLEN<br>DRQQTHMLDVMQDHFSRASSIIDELFQDRFFTREPQDTYHYL<br>PFSLPHRRPHFFFPKSRIVRSLMPFSPYEPLNFHAMFQPFLEMI<br>HEAQQAMDIHFHSPAFQHPPTEFIREGDDDRTVCREIRHNSTG<br>CLRMKDQCDKCREILSVDCSTNNPSQAKLRRELDESLQVAER<br>LTRKYNELLKSYQWKMLNTSSLLEQLNEQFNWVSRLANLTQ<br>GEDQYYLAAAAAAAATSDSDVPSGVTEVVVKLFDSDPITVT<br>VPVEVSRKNPKFMETVAEKALQEYRKKHREE**EPEA** |
| CLU-Ala$^{393-400}$-CT<br>(mt-15) | MMKTLLLFVGLLLTWESGQVLGDQTVSDNELQEMSNQGSK<br>YVNKEIQNAVNGVKQIKTLIEKTNEERKTLLSNLEEAKKKKE<br>DALNETRESETKLKELPGVCNETMMALWEECKPCLKQTCMK<br>FYARVCRSGSGLVGRQLEEFLNQSSPFYFWMNGDRIDSLLEN<br>DRQQTHMLDVMQDHFSRASSIIDELFQDRFFTREPQDTYHYL<br>PFSLPHRRPHFFFPKSRIVRSLMPFSPYEPLNFHAMFQPFLEMI<br>HEAQQAMDIHFHSPAFQHPPTEFIREGDDDRTVCREIRHNSTG<br>CLRMKDQCDKCREILSVDCSTNNPSQAKLRRELDESLQVAER<br>LTRKYNELLKSYQWKMLNTSSLLEQLNEQFNWVSRLANLTQ<br>GEDQYYLRVTTVASHAAAAAAAAGVTEVVVKLFDSDPITVT<br>VPVEVSRKNPKFMETVAEKALQEYRKKHREE**EPEA** |
| | MMKTLLLFVGLLLTWESGQVLGDQTVSDNELQEMSNQGSK<br>YVNKEIQNAVNGVKQIKTLIEKTNEERKTLLSNLEEAKKKKE<br>DALNETRESETKLKELPGVCNETMMALWEECKPCLKQTCMK<br>FYARVCRSGSGLVGRQLEEFLNQSSPFYFWMNGDRIDSLLEN |

| | |
|---|---|
| CLU-Ala$^{401-408}$-CT<br>(mt-16) | DRQQTHMLDVMQDHFSRASSIIDELFQDRFFTREPQDTYHYL<br>PFSLPHRRPHFFFPKSRIVRSLMPFSPYEPLNFHAMFQPFLEMI<br>HEAQQAMDIHFHSPAFQHPPTEFIREGDDDRTVCREIRHNSTG<br>CLRMKDQCDKCREILSVDCSTNNPSQAKLRRELDESLQVAER<br>LTRKYNELLKSYQWKMLNTSSLLEQLNEQFNWVSRLANLTQ<br>GEDQYYLRVTTVASHTSDSDVPSAAAAAAAALFDSDPITVTV<br>PVEVSRKNPKFMETVAEKALQEYRKKHREE**EPEA** |
| CLU-Ala$^{409-416}$-CT<br>(mt-17) | MMKTLLLFVGLLLTWESGQVLGDQTVSDNELQEMSNQGSK<br>YVNKEIQNAVNGVKQIKTLIEKTNEERKTLLSNLEEAKKKKE<br>DALNETRESETKLKELPGVCNETMMALWEECKPCLKQTCMK<br>FYARVCRSGSGLVGRQLEEFLNQSSPFYFWMNGDRIDSLLEN<br>DRQQTHMLDVMQDHFSRASSIIDELFQDRFFTREPQDTYHYL<br>PFSLPHRRPHFFFPKSRIVRSLMPFSPYEPLNFHAMFQPFLEMI<br>HEAQQAMDIHFHSPAFQHPPTEFIREGDDDRTVCREIRHNSTG<br>CLRMKDQCDKCREILSVDCSTNNPSQAKLRRELDESLQVAER<br>LTRKYNELLKSYQWKMLNTSSLLEQLNEQFNWVSRLANLTQ<br>GEDQYYLRVTTVASHTSDSDVPSGVTEVVVKAAAAAAAAVT<br>VPVEVSRKNPKFMETVAEKALQEYRKKHREE**EPEA** |
| CLU-Ala$^{417-424}$-CT<br>(mt-18) | MMKTLLLFVGLLLTWESGQVLGDQTVSDNELQEMSNQGSK<br>YVNKEIQNAVNGVKQIKTLIEKTNEERKTLLSNLEEAKKKKE<br>DALNETRESETKLKELPGVCNETMMALWEECKPCLKQTCMK<br>FYARVCRSGSGLVGRQLEEFLNQSSPFYFWMNGDRIDSLLEN<br>DRQQTHMLDVMQDHFSRASSIIDELFQDRFFTREPQDTYHYL<br>PFSLPHRRPHFFFPKSRIVRSLMPFSPYEPLNFHAMFQPFLEMI<br>HEAQQAMDIHFHSPAFQHPPTEFIREGDDDRTVCREIRHNSTG<br>CLRMKDQCDKCREILSVDCSTNNPSQAKLRRELDESLQVAER<br>LTRKYNELLKSYQWKMLNTSSLLEQLNEQFNWVSRLANLTQ<br>GEDQYYLRVTTVASHTSDSDVPSGVTEVVVKLFDSDPITAAA<br>AAAAARKNPKFMETVAEKALQEYRKKHREE**EPEA** |
| CLU-Ala$^{-425-432}$-CT<br>(mt-19) | MMKTLLLFVGLLLTWESGQVLGDQTVSDNELQEMSNQGSK<br>YVNKEIQNAVNGVKQIKTLIEKTNEERKTLLSNLEEAKKKKE<br>DALNETRESETKLKELPGVCNETMMALWEECKPCLKQTCMK<br>FYARVCRSGSGLVGRQLEEFLNQSSPFYFWMNGDRIDSLLEN<br>DRQQTHMLDVMQDHFSRASSIIDELFQDRFFTREPQDTYHYL<br>PFSLPHRRPHFFFPKSRIVRSLMPFSPYEPLNFHAMFQPFLEMI<br>HEAQQAMDIHFHSPAFQHPPTEFIREGDDDRTVCREIRHNSTG<br>CLRMKDQCDKCREILSVDCSTNNPSQAKLRRELDESLQVAER<br>LTRKYNELLKSYQWKMLNTSSLLEQLNEQFNWVSRLANLTQ<br>GEDQYYLRVTTVASHTSDSDVPSGVTEVVVKLFDSDPITVTVP<br>VEVSAAAAAAAATVAEKALQEYRKKHREE**EPEA** |
| | **MMKTLLLFVGLLLTWESGQVLG**DQTVSDNELQEMSNQGS<br>KYVNKEIQNAVNGVKQIKTLIEKTNEERKTLLSNLEEAKKKK<br>EDALNETRESETKLKELPGVCNETMMALWEECKPCLKQTCM |

| | |
|---|---|
| CLU-Ala$^{433-441}$-CT (mt-20) | KFYARVCRSGSGLVGRQLEEFLNQSSPFYFWMNGDRIDSLLE NDRQQTHMLDVMQDHFSRASSIIDELFQDRFFTREPQDTYHY LPFSLPHRRPHFFFPKSRIVRSLMPFSPYEPLNFHAMFQPFLEM IHEAQQAMDIHFHSPAFQHPPTEFIREGDDDRTVCREIRHNST GCLRMKDQCDKCREILSVDCSTNNPSQAKLRRELDESLQVAE RLTRKYNELLKSYQWKMLNTSSLLEQLNEQFNWVSRLANLT QGEDQYYLRVTTVASHTSDSDVPSGVTEVVVKLFDSDPITVT VPVEVSRKNPKFME**AAAAAAAAA**YRKKHREE**EPEA** |
| CLU-Ala$^{442-449}$-CT (mt-21) | MMKTLLLFVGLLLTWESGQVLGDQTVSDNELQEMSNQGSK YVNKEIQNAVNGVKQIKTLIEKTNEERKTLLSNLEEAKKKKE DALNETRESETKLKELPGVCNETMMALWEECKPCLKQTCMK FYARVCRSGSGLVGRQLEEFLNQSSPFYFWMNGDRIDSLLEN DRQQTHMLDVMQDHFSRASSIIDELFQDRFFTREPQDTYHYL PFSLPHRRPHFFFPKSRIVRSLMPFSPYEPLNFHAMFQPFLEMI HEAQQAMDIHFHSPAFQHPPTEFIREGDDDRTVCREIRHNSTG CLRMKDQCDKCREILSVDCSTNNPSQAKLRRELDESLQVAER LTRKYNELLKSYQWKMLNTSSLLEQLNEQFNWVSRLANLTQ GEDQYYLRVTTVASHTSDSDVPSGVTEVVVKLFDSDPITVTVP VEVSRKNPKFMETVAEKALQE**AAAAAAAA****EPEA** |

**Note:**

1. Key:

   MMKTLLLFVGLLLTWESGQVLG: Signal peptide sequence
   **AAAAAAAA**: Alanine stretch substitutions
   **EPEA**: C-tag sequence
2. The superscript numbering refers to the amino acid residue number in mature human CLU.
3. Identification codes for alanine substituted mutants (e.g. mt-1) are shown in parentheses under the full name.

**Table S4 Yield of purified CLU-CT proteins transiently expressed in MEXi 293E cells.**

| **Protein** | **Yield (mg/L)** |
|---|---|
| $^{wt}$CLU-CT (wt) | 30 |
| CLU-ala$^{369-376}$-CT (mt-12) | 20 |
| CLU-ala$^{377-384}$-CT (mt-13) | 4 |
| CLU-ala$^{442-449}$-CT (mt-21) | 7 |

**Table S5 Amino acid sequence of dNbs generated against specific CLU epitopes and control dNb.**

| dNb | CLU target epitope (residues) | Amino acid sequence of the complete dNbs including binding region (in red) and C-tag (underlined) | $A_{0.1}$% (Extinction Coefficient) |
|---|---|---|---|
| $dNb^{369-376}$ | VSRLANLT (369-376) | MRGSHHHHHHGMASMTGGQQMGRDLYDDDDKDPKLEVQLVESGGGLVQPGGSLRLSCAASGFNIKDTYIGWVRRAPGKGEEWVASIYPTNGYTRYADSVKGRFTISADTSKNTAYLQMNSLRAEDTAVYYCAAGSKLAVHSYGEEEAAAWGQGTLVTVSSGTEPEA | 2.080 |
| $dNb^{442-449}$ | YRKKHREE (442-449) | MRGSHHHHHHGMASMTGGQQMGRDLYDDDDKDPKLEVQLVESGGGLVQPGGSLRLSCAASGFNIKDTYIGWVRRAPGKGEEWVASIYPTNGYTRYADSVKGRFTISADTSKNTAYLQMNSLRAEDTAVYYCAAGSRTVELEVEEEAAAWGQGTLVTVSSGTEPEA | 1.976 |
| $dNb^{neg}$ | none | MRGSHHHHHHGMASMTGGQQMGRDLYDDDDKDPKLEVQLVESGGGLVQPGGSLRLSCAASGFNIKDTYIGWVRRAPGKGEEWVASIYPTNGYTRYADSVKGRFTISADTSKNTAYLQMNSLRAEDTAVYYCAAGSGASASAGAEEEAAAWGQGTLVTVSSGTEPEA | 1.749 |

**Table S6 CLU ligand binding sites predicted using FTMove [1]**

| CLU residues (no. residues) | Confidence score (max 1) |
|---|---|
| S370-Y382 (12) | 0.999 |
| S228-A242 (14) | 0.999 |
| N157-L172 (15) | 0.993 |
| N363-W368 (6) | 0.872 |
